# NLRP3- and AIM2-autonomy in a mouse model of MSU crystal-induced acute inflammation *in vivo* highlights imiquimod-dependent targeting of *Il-1β* expression as relevant therapy for gout patients

**DOI:** 10.1101/772756

**Authors:** Alexandre Mariotte, Aurore De Cauwer, Chrystelle Po, Chérine Abou-Faycal, Angélique Pichot, Nicodème Paul, Ismael Aouadi, Raphael Carapito, Benoit Frisch, Cécile Macquin, Emmanuel Chatelus, Jean Sibilia, Jean-Paul Armspach, Seiamak Bahram, Philippe Georgel

## Abstract

The role of Monosodium Urate (MSU) crystals in gout pathophysiology is well described, as is the major impact of IL-1β in the inflammatory reaction that constitutes the hallmark of the disease. However, despite the discovery of the NLRP3 inflammasome and its role as a Pattern Recognition Receptor linking the detection of a danger signal (MSU) to IL-1β secretion *in vitro*, the precise mechanisms leading to joint inflammation in gout patients are still poorly understood. Here, we provide an extensive clinical, biological and molecular characterization of the acute uratic inflammation mouse model induced by subcutaneous injection of MSU crystals, which accurately mimics human gout. Our work reveals several key features of MSU-dependent inflammation and identifies novel therapeutic opportunities, among which the use of topical application of imiquimod to promote interferon-dependent anti-inflammatory action maybe relevant.

## Introduction

Gout is a painful inflammatory arthritis, which exhibits an increasing prevalence worldwide, thereby representing a public health issue (1). While systemic hyperuricemia leading to the accumulation of Monosodium Urate (MSU) crystals is a well-established etiological cause of the disease, the precise understanding of the molecular and cellular mechanisms driving joint inflammation is no yet achieved. Genome-wide association studies (GWAS) performed in Japanese patients and controls evidenced variants in genes encoding proteins involved in urate transport and metabolism (2), and the prominent role of urate transporters was confirmed in various populations (3). More recently, a GWAS realized on Taiwanese patients revealed 36 loci associated with gout, among which rs2231142 in the *ABCG2* gene exhibited the strongest linkage (4). Of note, *ABCG2* knockdown in endothelial cells induces increased secretion of the neutrophil chemoattractant IL-8, which might provide some mechanistic insights into gout pathophysiology. Surprisingly, none of these pan-genomic studies - some of which were performed in considerably large cohorts (e.g. 70,000 patients (5)) - revealed any linkage with genes encoding components of the NLRP3 inflammasome. Gain-of-function mutations in *NLRP3* are responsible for cryopyrin-associated autoinflammatory syndrome (CAPS) characterized by systemic inflammation (fever and urticaria-like rash) not evocative of gout, although arthralgia and periarticular swelling can occur (6). This appears in sharp contrast with the standard model which stipulates that MSU crystals trigger activation of the NLRP3 inflammasome, ultimately leading to IL-1β secretion (7). Indeed, a large body of evidence indicates that *NLRP3* deficiency prevents IL-1β secretion by murine macrophages following MSU crystals stimulation (8). Human mononuclear cells also respond to MSU crystals which, in synergy with lipopolysaccharide (LPS), enhance IL-1β production (9). However, these experiments require the presence of LPS as a “priming signal” necessary for optimal IL-1β release, a component which is likely absent in gouty patients. Recently, neutrophils, which are abundant in the inflamed synovial fluids of acute patients, also appeared as major players in gouty arthritis (10). Moreover, these cells are also involved during the resolutive phase through NETosis (11). In the present work, we sought to gain additional insights into cellular and molecular interactions driving gouty arthritis and for this, we performed subcutaneous (s.c) injections of MSU crystals in the hind paws of mice, a model that accurately mimics the human disease, both clinically as well as pharmacologically. In this setting, we observed that both the NLRP3 and AIM2 inflammasomes are, at best, only partially required for the full development of the disease *in vivo*. This likely indicates that multiple pathways may act redundantly to promote the release of bioactive IL-1β, and highlights the necessity to target *Il-1β* gene expression for efficient therapeutic purposes. We next analyzed this complex MSU-induced inflammatory transcriptional response *in vivo* by genome-wide RNAseq and observed a marked overexpression of genes involved in granulocyte adhesion and diapedesis (e.g. *Itgam*, chemoattractants such as *Ccl3* or *Cxcl3)*, as well as genes participating in the NLRP3 inflammasome pathway (*Nlrp3, Il-1β)*. Interestingly, we also noticed *Tlr-1* and *-6* over expression, pointing to a potential involvement of endogenous lipids as the - still elusive - signal I in gouty patients. Similarly to our previous observation in murine models of arthritis (12), we next demonstrated that type I interferon anti-inflammatory effects can be mobilized through topical application of the TLR7 agonist imiquimod on inflamed paws in our gout model. The resulting lower expression of inflammasome-related genes, including *Il-1β* and neutrophils recruitment players, correlated with a drastic reduction of the MSU-induced inflammation, both at the clinical and biological levels. Furthermore, our transcriptomic analyses uncovered previously unsuspected players (*RUNX3*) accounting for this type I IFN-dependent anti-inflammatory response. Altogether, our results offer novel therapeutic opportunities, particularly for gout patients with kidney dysfunctions in which colchicine, the standard of care, might be inadvisable.

## Results

### Subcutaneous injection of MSU crystals in mice recapitulates human gouty arthritis

To investigate molecular and cellular relationships that participate in gouty arthritis, we used a mouse model in which MSU crystals are injected subcutaneously (s.c) on the dorsal face of hind paws (11, 13). As seen in Suppl. Fig. 1 A-B, mice subjected to this protocol develop an oedema, which can reach the ankle area within 24h (acute phase). However, the inflammation gradually decreases after seven days (168h) and MSU crystals can be visible under the skin as tophi-like structures. Magnetic resonance imaging (MRI, panel C) of MSU-injected paws revealed tenosynovitis, subcutaneous oedema and crystal deposits which were absent in PBS-injected contralateral paws. These manifestations correlated with peak production of mature (17 kDa) IL-1β at 24h post MSU crystal injection (p.i) (Panel D). Of note, paw thickness, as well as inflammatory score (which depends on both redness and swelling) were significantly reduced when mice were treated with colchicine, the standard of care for human gout (Suppl. Fig. 2A-D). Increased body temperature, reflecting the systemic effects of MSU injection, was also impacted by colchicine therapy. Furthermore, this model was responsive to the IL-1 receptor antagonist Anakinra (Suppl. Fig. 2E-F), at least during the initial phase of the disease (which probably reflects its limited half-life (14)), but not to the TNF-α blocker Etanercept (Suppl. Fig. 2G-H). Because of this partial antibody-mediated IL-1β inhibition, we performed MSU-induced acute inflammation in *Il-1β* KO animals and observed a major reduction of paw swelling and clinical scores (Suppl. Fig 3A-B), concomitantly with a drastic decrease of tissue IL-6 and Myeloperoxydase (MPO, Suppl. Fig. 3 C-E). On the contrary, MSU-injected *Il-1α* KO mice exhibited only limited reduction in tissue IL-1β and MPO, with clinical symptoms of joint inflammation similar to wild types. Altogether, these data ascertained the s.c injection of MSU crystals in mice as an acute uratic inflammation model, highly dependent on massive IL-1β secretion, which appears to mimic human gout.

### The PKD - NLRP3 axis is dispensable for gouty arthritis in mice

NLRP3 is described as a sensor of MSU crystals in macrophages (8) and considered a major player in the development of gout in various *in vivo* models (15, 16). However, this dogma regarding the central role of NLRP3 in gout has been challenged (17, 18), although IL-1β remains a crucial cytokine in the development of the disease. In our model, *Nlrp3*-deficient mice exhibited similar joint swelling and clinical score compared with littermate controls (Fig. 1 A, B), despite showing reduced fever 24 h after MSU challenge (Panel C). We therefore checked IL-1β expression in our model. Both pro and mature forms were similarly detected by Western blot in the paws of mutant and wild type mice during the acute phase (24h) after MSU injection (Fig. 1D). Accordingly, quantification of IL-1β by tissue ELISA (which detects both pro and mature forms) showed equivalent amounts in paws from MSU-stimulated *Nlrp3* mutants and controls 24h post MSU injection (Fig. 1E). Finally, myeloperoxidase quantification revealed similar neutrophil infiltration in the paws of *Nlrp3* ^-/-^ and wild type animals following MSU injection (Fig. 1F). Similar results were obtained following pharmacological (using CRT0066101) inhibition of Protein Kinase D (PKD), an essential component enabling NLRP3 inflammasome activation (19) (Suppl. Fig. 4 A-E). To check the validity of the genetic and pharmacological inhibition of NLRP3, we performed several *in vitro* experiments. In line with published reports (8, 19), macrophages and neutrophils harvested from *Nlrp3* ^-/-^ mice or wild type cells treated with CRT0066101 failed to secrete IL-1β upon MSU or ATP stimulation following LPS priming (Suppl. Fig 5 A, B), while TNF-α and IL-6 secretion remained unaffected (Suppl. Fig. 5C). We also noted that neutrophils, as opposed to macrophages, appeared unresponsive to a corpuscular trigger (e.g. MSU crystals) compared to a soluble one (such as ATP) for NLRP3-dependent IL-1β production, a feature that has been observed by others (20).

**Figure 1.**
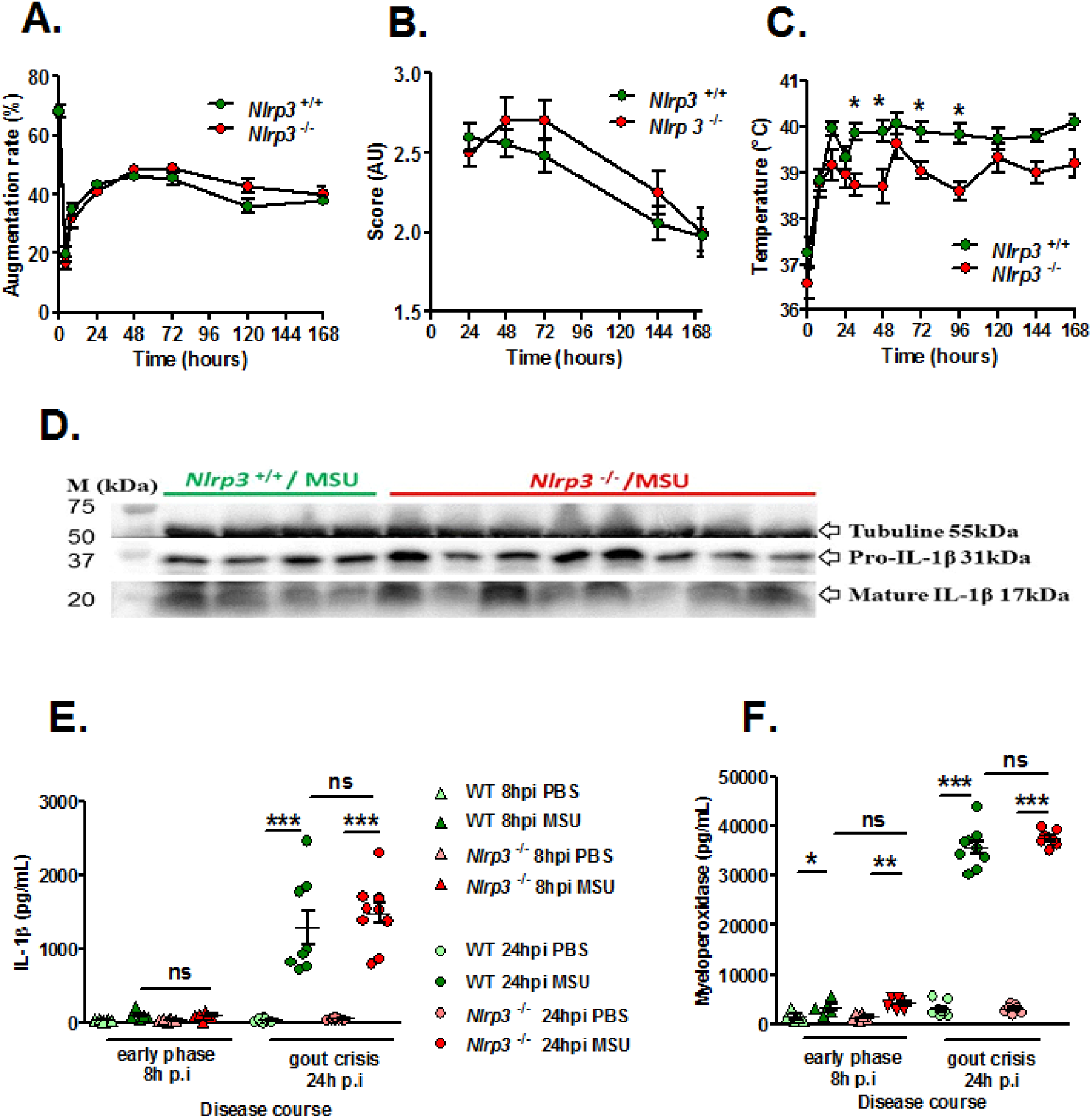
The NLRP3 inflammasome is not required for IL-1β maturation and consecutive inflammation in the subcutaneous acute uratic inflammation model. **A-F.** *Nlrp3*^*+/+*^ and *Nlrp3*^*-/-*^ mice were submitted to an acute uratic inflammation experiment. **A.** Paw swelling, **B.** Clinical scores and **C.** Body temperature were recorded over a 7-days-long period (n = 11 to 13 in each group). **D-F.** *In situ* (paws) analysis of IL-1β production/maturation and MPO quantification. **D.** 24 hours post-injection, paws were harvested and a western blot analysis was realized on 75µg of paw extracts to visualize pro-IL-1β (31kDa) and its cleaved, mature form (17kDa); Tubulin was used a loading control (n = 4 *Nlrp3*^*+/+*^ and n=8 *Nlrp3*^*-/-*^). **E.** IL-1β and **F.** myeloperoxidase (MPO) were quantified by ELISA in paw extracts obtained at the early phase (8h post-injection, n = 5 to 6) and the peak (24hpi, n = 8 to 10). All data are representative of three independent experiments except for clinical observation (five independent experiments). Symbols represent individual mice, horizontal lines and bars correspond to mean +/- SEM; Results were analyzed with a two-tailed Mann-Whitney test, * = p<0.05, ** = p < 0.01, *** = p < 0.001, ns = not significant.

Furthermore, we performed intraperitoneal injection of MSU and observed that this inflammation model is also *Nlrp3*-independent (Suppl. Fig. 6), as assessed by similar number of infiltrating neutrophils and macrophages in the peritoneal cavity of wild type and mutant mice 6h upon MSU injection (panels B-D) and the comparable amount of IL-1β in the peritoneal fluid (panel E). Our data, while confirming that the NLRP3 inflammasome is mandatory in isolated cells (macrophages, neutrophils) to respond to MSU, also suggest that this inflammasome is dispensable for *in vivo* joint inflammation following s.c injection of urate crystals. To gain more insight into the early response (acute phase) in the paws following MSU s.c injection, we performed a genome-wide RNAseq analysis to compare MSU-injected paws to those receiving PBS as control at 24 hours post injection (hpi). As seen on the volcano plot shown in Suppl. Fig. 7A, a massive induction of gene transcription occurred following s.c MSU injection and the heatmap (panel B) of the significantly (p<0.001) 405 differentially expressed genes shows convincing unsupervised clustering of the Control (PBS) and MSU-injected samples. The heatmap of the top 100 genes (panel C) highlighted several features, among which the overexpression of *Il-1b* validated our data. Interestingly, *Tlr1* and *Tlr6* overexpression was also evidenced, pointing to a role for endogenous lipids as possible inducers of the signal I activating NF-κB signaling (18). Such a signal is generally provided by LPS during experimental *in vitro* models of inflammasome activation by MSU crystals, but its physiological relevance is even more questionable in light of our data showing that *Tlr4* expression remained unaffected in MSU-challenged mice. Finally, *Ccl3* strong overexpression might also account for the high fever observed upon MSU crystals injection (21). As expected, Gene Ontology (GO) terms analysis (panel D) revealed an enrichment in immune responses/inflammatory responses upon MSU injection. A comparison of 211 differentially expressed genes (showing a highly statistically significant difference between PBS- and MSU-injected paws p<0.0001) using Ingenuity Pathways Analysis (IPA, Qiagen) and Reactome (https://reactome.org/) revealed a significant (p=2.53E-24) enrichment in genes required for granulocyte adhesion, diapedesis and degranulation (see IPA Network in Suppl. Fig. S8A). Finally, we used the ImmQuant tool (22) to extrapolate the immune cell composition from our RNAseq data. In line with enhanced MPO detection (Suppl. Fig. S3 and Fig. 1) and overexpression of neutrophil chemoattractants (*Cxcl1* and *Cxcl2*), MSU s.c injection triggered granulocyte and monocyte recruitment (Suppl. Fig. S9, panels A and D).

### MSU crystal-induced arthritis can develop autonomously from both NLRP3 and AIM2 inflammasomes

Neutrophilic infiltration and subsequent NETosis, a feature of inflamed joint in gout patients (23), exposes self-DNA, which is suspected - once phagocytozed - to induce AIM2-dependent IL-1β secretion (reviewed in (24)). This prompted us to investigate the potential role of this alternative inflammasome in our model of acute experimental gout. Like *Nlrp3* mutants, mice lacking *Aim2* displayed inflammatory responses similar to those observed in controls following MSU crystal injection (Fig. 2 A-C). Next, to explore a possible redundancy between the two inflammasomes, we produced *Nlrp3* ^-/-^; *Aim* 2 ^-/-^ mice. As seen in panels D-F, double mutant mice exhibited slightly reduced clinical responses (with differences in the area under the curve of the measure of the paw swelling reaching statistical significance between wild type and double KO mice) upon MSU crystal s.c injection. However, quantification of IL-1β and myeloperoxidase (MPO) by tissue ELISA 24h upon MSU challenge, as well as seric IL-1β dosage (12 and 24 h after MSU injection, Fig 2. G-H), confirmed that MSU-dependent rise in these biological inflammatory markers can occur independently from these two inflammasomes. Similar observations could be made from the analysis of *Caspase 1/11*-deleted mice (*Casp 1*^*-/-*^, *11* ^*-/-*^) in which MSU injection induced a weak reduction of the inflammatory symptoms, albeit without showing any effect on IL-1β production (Suppl. Fig. S10).

**Figure 2.**
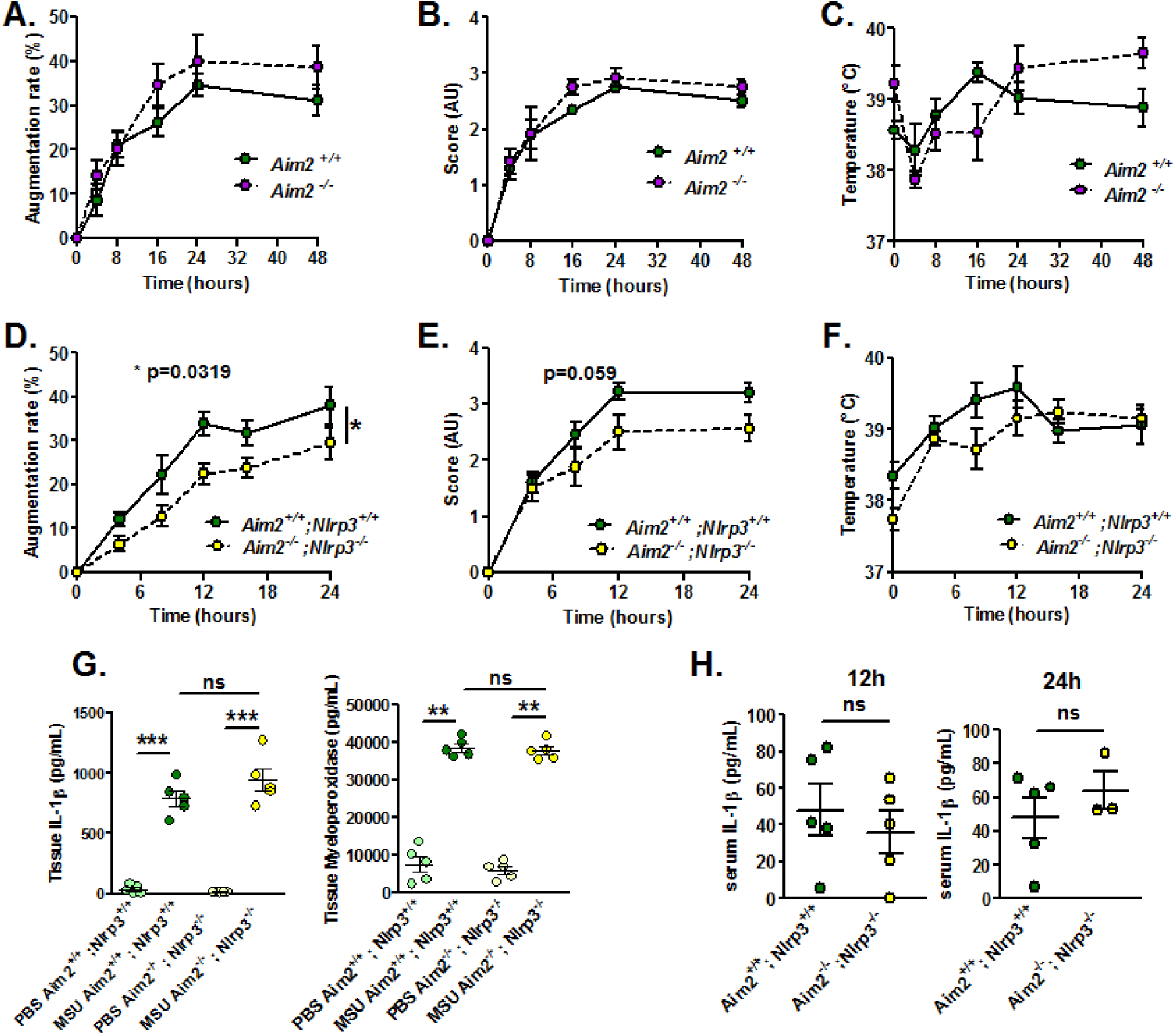
*Aim2* or *Aim2/Nlrp3* combined deficiencies do not impact on acute uratic inflammation development and severity. **A-C.** Acute uratic inflammation was induced in *Aim2*^*+/+*^ and *Aim2*^*-/-*^ animals (n = 3 to 6). **A.** Paw swelling, **B.** Clinical scores and **C.** Body temperature. **D-E.** *Aim2*^*-/-*^; *Nlrp3* ^*-/-*^ (double knock out) mice (n = 8) and *Aim2*^*+/+*^; *Nlrp3*^*+/+*^ (n = 10) were subjected to an acute uratic inflammation experiment. **D.** Paw swelling and **E.** Clinical scores and **F.** body temperature measures, all recorded over a 24h period. **G.** IL-1β and MPO quantified by ELISA in paws extracts collected at 24hpi, n = 5 in each group**. H.** IL-1β and MPO quantified by ELISA in serum collected 12hpi (left) and 24hpi (right) (n = 3 to 5). Symbols represent individual mice; green dots correspond to WT mice (littermate controls), purple dots to *Aim2*^*-/-*^ mice and yellow dots to *Aim2*^*-/-*^; *Nlrp3*^*-/-*^ mice; light colored dots represent PBS paws and bright colored dots the MSU-injected paws. Horizontal lines and bars correspond to mean +/- SEM; Results were analyzed with a two-tailed Mann-Whitney test, * = p<0.05, ** = p < 0.01, *** = p < 0.01, ns = not significant. In graphs D. and E. the area under curve (AUC) was determined and analyzed by a Mann-Whitney test.

Since *Nlrp3*-deleted macrophages can efficiently (and even more than controls) respond to transfected poly A:T, a classical inducer of the AIM2 inflammasome (Fig. 3A), we tested the existence of a possible collaboration between neutrophils, in which MSU crystals poorly induce IL-1β secretion (Suppl. Fig. 5) but efficiently trigger DNA release through NETosis (23), and macrophages, major producers of IL-1β once triggered by AIM2-dependent signals (such as poly dA:dT). For this, we performed co-cultures of both cell types isolated from wild type or *Nlrp3* ^-/-^; *Aim2* ^-/-^ double knock-out (dKO) animals. As seen in Fig. 3B, IL-1β release by control macrophages did not significantly differed whether they are cultured with neutrophils with or without MSU. This indicates that NETosis triggered by MSU ((11, 23) and our own observation) did not affect IL-1β release, thus suggesting that DNA exposure is not essential in this process and cannot provide the necessary signal I driving *Il-1β* transcription.

**Figure 3.**
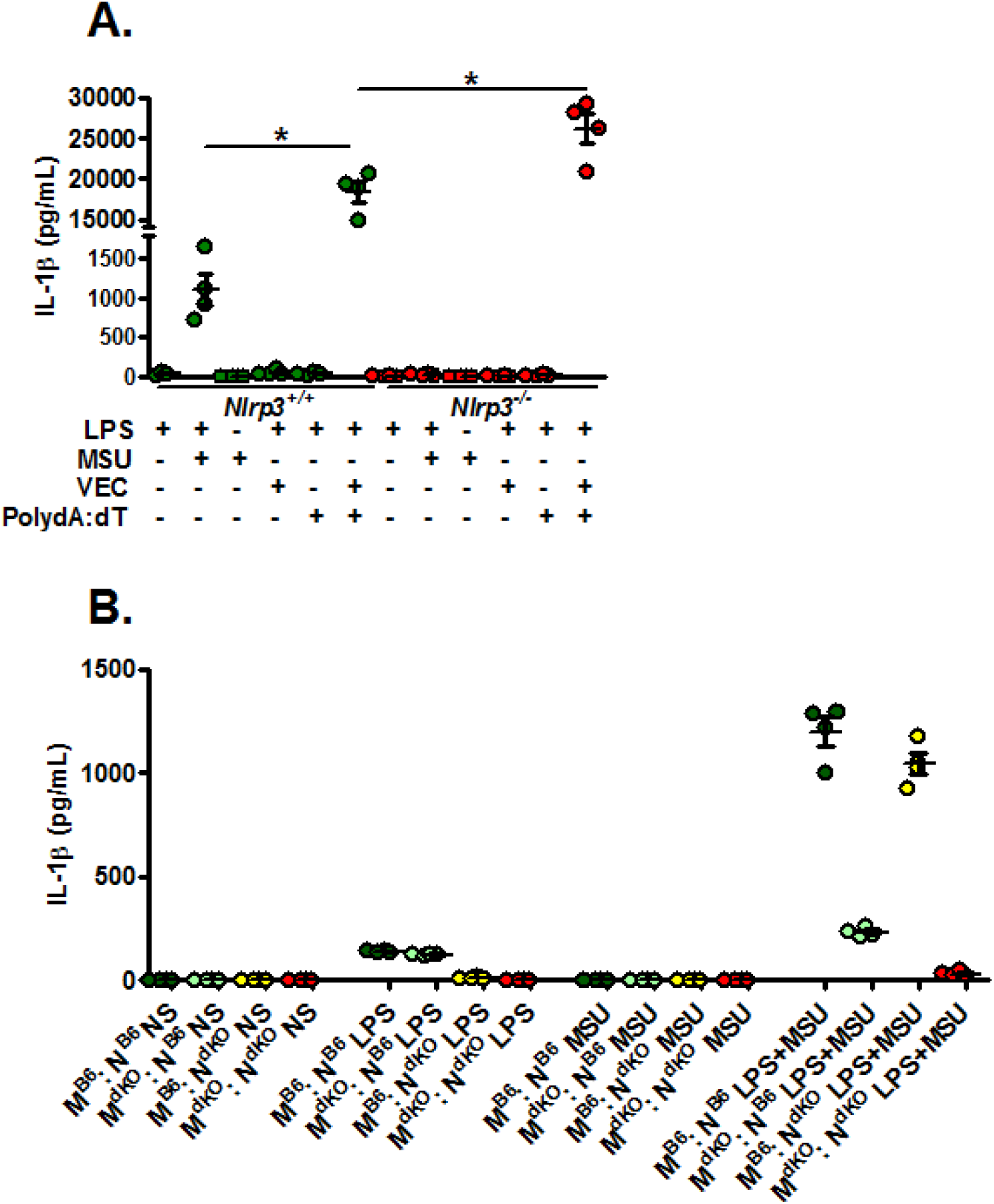
Activated (netotic) neutrophils fail to promote IL-1β secretion by macrophages. **A.** *Nlrp3*^*+/+*^ and *Nlrp3*^*-/-*^ peritoneal macrophages were seeded at 5×105 cells/well and stimulated with LPS only with LPS and MSU, LPS and lipofectamin (VEC), LPS and poly dA:dT (0,5µg/mL) or LPS and lipofectamin complexed with poly dA:dT (0,5µg/mL). Green dots represent *Nlrp3*^*+/+*^ macrophages, red dots *Nlrp3*^*-/-*^ cells. **B.** WT (*C57Bl/6* or *B6*) or *Aim2*^*-/-*^; *Nlrp3*^*-/-*^ (double knock out: dKO) peritoneal macrophages (M) were plated at 5×10^5^ cells/well together with 1×10^6^ neutrophils (N) of both genotype making combinations (M B6 : N B6, dark green; M dKO : N B6, light green; M B6 : N dKO, yellow; M dKO : N dKO, red). Cells were left untreated (NS) or treated with LPS only (1µg/mL for 27h), MSU crystals only (250µg/mL) for 24h or both (LPS alone for 3 hours and then MSU for 24h). Cells were isolated from n=4 mice. Horizontal lines and bars correspond to mean +/- SEM; Results were analyzed with a two-tailed Mann-Whitney test, * = p<0.05, ns = not significant.

### Imiquimod downmodulates *Il-1β* gene expression and protects from experimental gouty arthritis

We previously reported the therapeutic effects of topical imiquimod, a known TLR7-dependent inducer of type I IFN responses in mouse models of rheumatoid arthritis (12) and its effect on neutrophil recruitment. Therefore, we questioned the potential benefits of this treatment in our present gout model, which is characterized by a transcriptomic signature revealing MSU-induced genes involved in granulocyte diapedesis and activation. Except for body temperature, topical application of imiquimod caused a marked reduction of the clinical parameters (paw swelling, redness, Fig. 4 A-E) associated with s.c injection of MSU. Tissue ELISA revealed that imiquimod specifically reduced IL-1β and IL-6 secretion in the treated paws (Panels F-G), whereas neutrophil infiltration (visualized by MPO quantification, Panel H) and TNF-α production (panel I) remained unaffected. RTqPCR and western blot analysis further documented the negative effect of imiquimod on the transcription of the *Il-1β* and *Il-6* genes (Panels J-K) and IL-1β pro-form expression (Panel L). Finally, we used a state-of-the-art imaging approach (MRI, Fig. 5) similar to that which is currently applied to monitor patients suffering various arthritides, including gout, to quantify the beneficial effects of imiquimod. This enabled us to better demonstrate the significant reduction of oedema formation and crystal deposit in imiquimod-treated paws. MRI also enabled us to visualize the reduction of tenosynovitis in imiquimod-treated animals (Suppl. Fig. S11).

**Figure 4.**
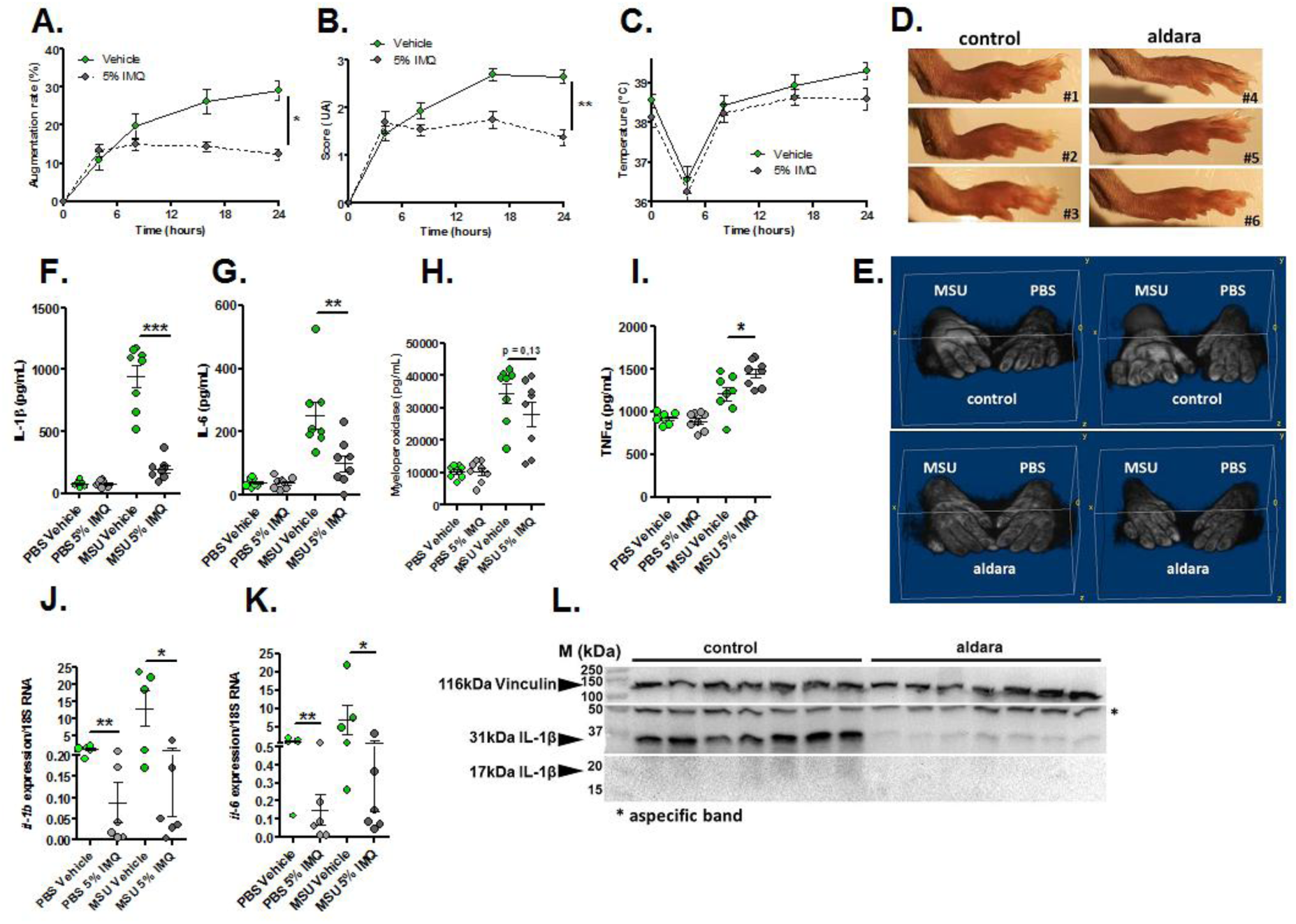
Topical application of an imiquimod-containing cream (ALDARA) alleviates acute uratic inflammation by down-modulating *Il-1b* and *Il-6* expression. **A-E.** C57BL/6J mice were submitted to an acute uratic inflammation experiment and immediately treated with either topical imiquimod (ALDARA® commercial cream) or a control cream. Gouty crisis was followed for 24 hours (n =8 mice per group, data are representative of 3 independent experiments). **A.** Paw swelling, **B.** Clinical scores, **C.** Body temperature, **D.** representative pictures of the paws (16hpi) and **E.** volumetric representation of 2 pairs of control cream-treated paws (upper panel), imiquimod-treated paws (lower panel) obtained after Magnetic Resonance Imaging (MRI) and image reconstruction with 3D viewer. **F.-I.** Paws were collected at 24hpi and protein extracts were analyzed by ELISA for IL-1β **(F.)**, IL-6 **(G.)**, MPO **(H.)** and TNFα **(I.). J.-K.** Similar to **F.-I.**, paws were collected at 24hpi and analyzed by RT-qPCR analysis for *Il-1β* **(J.)** and *Il-6* expression **(K.). L.** Proform and mature IL-1β production was assessed by western blot of paw extracts. Vinculin was used as loading control. Green dots represent control cream-treated mice, grey dots the topical imiquimod-treated animals; light colors correspond to PBS paws, dark colors to MSU paws. Symbols represent individual mice, horizontal lines and bars correspond to mean +/- SEM; Results were analyzed with a two-tailed Mann-Whitney test, * = p<0.05; ** = p < 0.01; *** = p < 0.001, ns = not significant. In graphs A and B, the area under the curve (AUC) was calculated and analyzed with a Mann-Whitney test.

**Figure 5.**
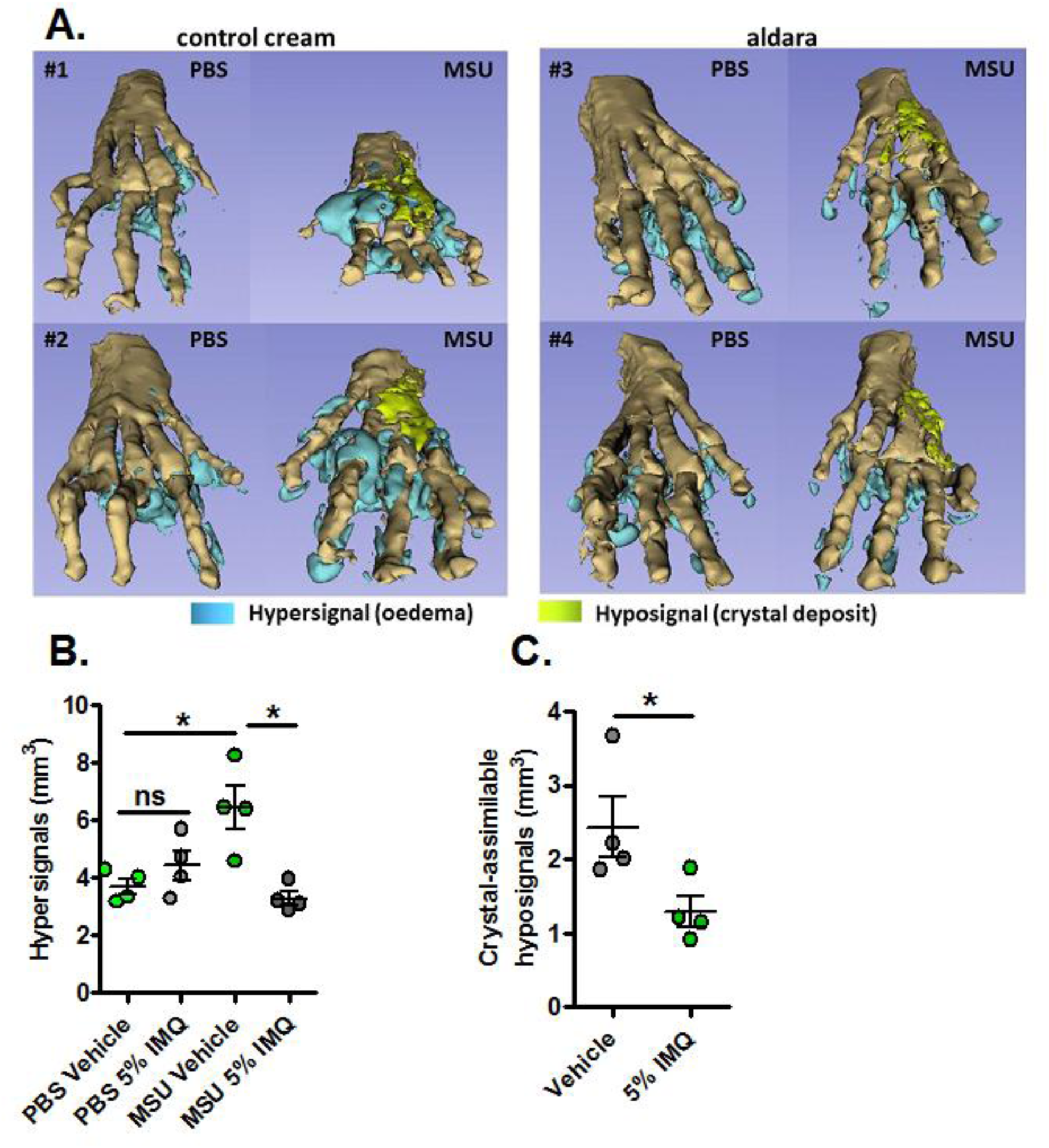
MRI reveals major dampening of MSU crystal-induced oedema formation upon topical imiquimod therapy. **A-C.** Acute uratic inflammation was induced and topical imiquimod or control cream were immediately applied on the paws of mice (n=4 per group). 24hpi, animals were euthanized and subjected to Magnetic Resonance Imaging (MRI). After acquisition of MRI sequences (T2, fat suppression), pictures were computed and reconstructions were made by 1) automated segmentation and 2) reconstruction of hyper-signal volumes (oedemas), hypo-signals volumes (MSU crystal-associated volume) and bones with the help of the 3D Slicer software. **A.** 3D reconstructions of 4 pairs of paws (control cream-treated, left; imiquimod-treated, right). **B.** Quantification of the hyper-signal volume. **C.** Quantification of the crystal-assimilable hypo-signal volumes. Green dots represent control cream-treated mice, grey dots miquimod-treated animals, light colors correspond to PBS paws, dark colors to MSU-injected paws. Symbols represent individual mice, horizontal lines and bars correspond to mean +/- SEM; Results were analyzed with a two-tailed Mann-Whitney test, * = p<0.05, ns = not significant.

Altogether, our data support the notion that the protective effect of Imiquimod-dependent TLR7 activation involves negative transcriptional control of genes encoding major pro-inflammatory cytokines, such as IL-1β and IL-6. To better describe, at the molecular level, the pathways that are modulated by imiquimod, we performed genome-wide transcriptomic on MSU-inflamed paws treated or not with imiquimod at 24 hpi. As a control, non-inflamed (PBS-injected) paws were also treated by topical application of imiquimod. In agreement with the reported role of imiquimod on its cognate receptor (TLR7), we identified a transcriptomic signature characterized by an overexpression of Interferon-stimulated genes (ISGs, such as *Oas1* and *2, Mx1, Isg15*; see Suppl. Fig. S12) in control mice (PBS-injected, imiquimod-treated). By contrast the combination MSU injection plus imiquimod induced an important down modulation of gene transcription (Suppl. Fig. S13A). Among the genes exhibiting highly significant transcriptional repression, *Il-1β*, as well as inflammasome components (e.g *Nlrp3*) appeared of particular importance (Suppl. Fig. S13C) and confirmed our RT-qPCR data (Fig. 4, panels F, J and L). Furthermore, reduction of *Cxcl2* expression is likely an important event contributing to the diminution of the neutrophilic infiltrate seen by ImmQuant analysis (Suppl. Fig. S9). Interestingly, an increased skin dendritic cell signature also seemed to take part in the anti-inflammatory effect of Imiquimod in the MSU/imiquimod combination, although this intriguing observation requires additional confirmation. Finally, MSU *vs* MSU/imiquimod comparative RNAseq analysis by IPA identified interconnected type I IFN-dependent genes. Interestingly, most genes involved in granulocyte adhesion and diapedesis, which were upregulated upon MSU stimulation, exhibited down-modulation in the samples from imiquimod-treated paws (see Suppl. Fig. S8B). Of note, these genes are connected in a network (predicted by IPA) in which SYK/AKT fill a central position (Suppl. Fig. S14). Although IL-1β repression by type I IFN has been previously described (25) and attributed to signaling through the IL-10 / STAT3 pathway (26), our dataset does not clearly point to such connection. Rather, reduced *Tlr6* expression and downstream NF-κB signaling is supported by the genes network built by IPA to account for diminished *Il-1b* expression. Unexpectedly, Reactome (https://reactome.org/PathwayBrowser/#/) identified a RUNX3 pathway in the imiquimod-treated samples, which may also support low *Il-1b* expression. Indeed, RUNX3, which has been described as a transcriptional regulator of *Il-1b* (27) is significantly (p=2,92991E-09) downmodulated by imiquimod. Altogether, our data suggest that the therapeutic properties of imiquimod in our experimental setting are mediated by the ability of this compound to impair two major hallmarks of inflammation: neutrophil recruitment and/or activation and IL-1β production.

Because our data point to a possible involvement or TLR-dependent signaling (possibly TLR1, 2 and 6) in our model of gouty arthritis, we tested the impact of MSU injection in *MyD88-* and *Il-1R1*-deficient animals. We observed an almost complete absence of inflammatory reactions (as judged by paw swelling and clinical score) in both *MyD88* and *Il-1R1* KO mice (Fig. 6A-B). However, we noticed that *Il-1R1* KO animals exhibited a significant residual expression of inflammatory markers (IL-1β, IL-6 and MPO), which were nearly undetectable in *MyD88*-depleted mice (Panels C-E). Because MYD88 is an adapter used downstream of both IL-1R1- and TLR-dependent signaling pathways (28), this observation strongly suggest that TLRs are indeed essential sensors driving NF-κB-dependent *Il-1β* transcription in response to MSU injection *in vivo*.

**Figure 6.**
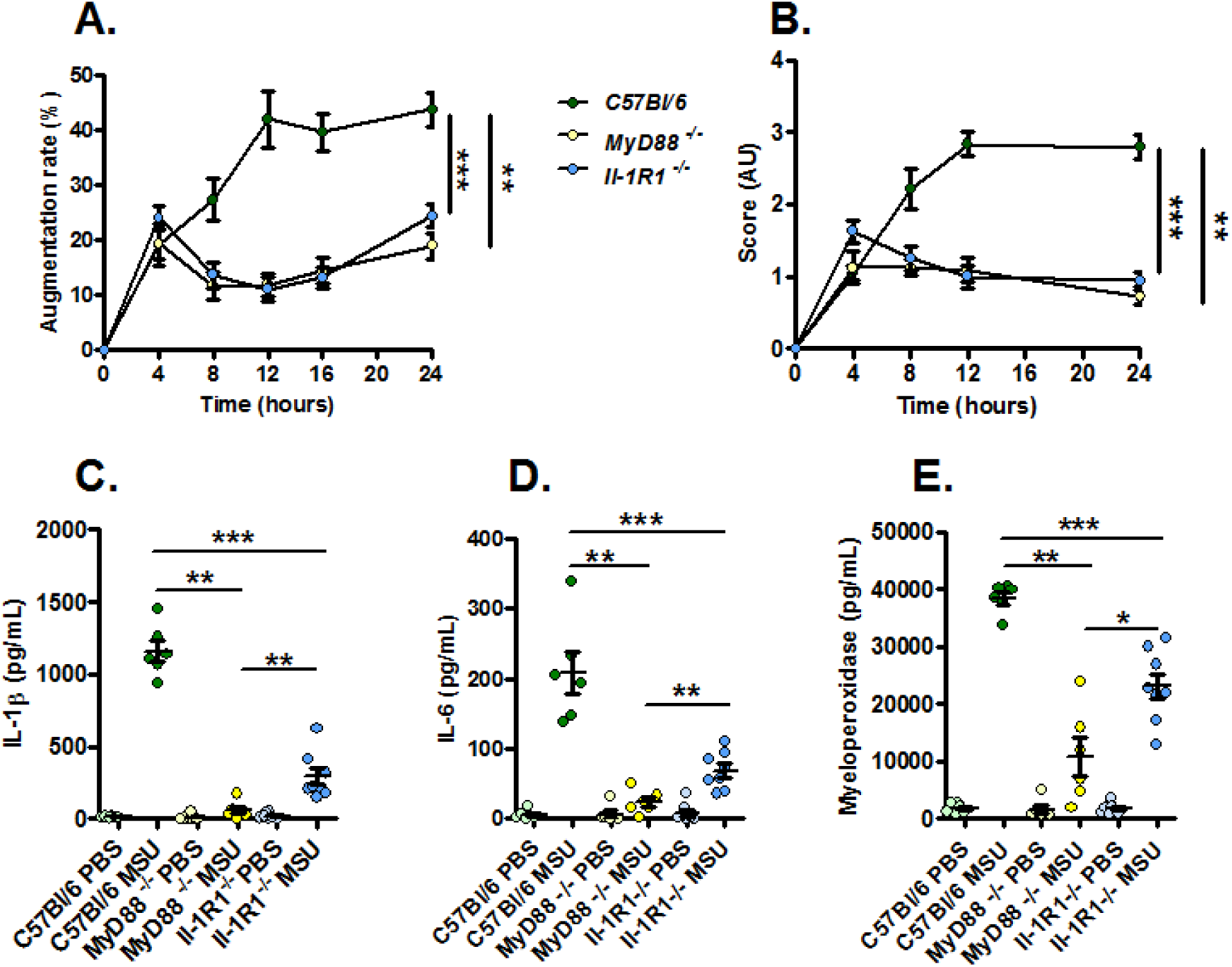
Increased resistance to MSU-induced inflammation in *Myd88* KO mice compared to *Il-1R1* KO animals. *C57Bl/6* (n=6, green dots), *Myd88* KO (n=6, yellow dots) and *Il-1R1* KO (n=8, blue dots) mice were submitted to an acute uratic inflammation experiment and clinical observations were recorded over a 24h period. **A.** Paw thickness, **B.** Clinical score. **C.** IL-1β, **D.** IL-6 and **F.** myeloperoxidase (MPO) were quantified by ELISA in protein extracts obtained from control paws 24h post PBS injection or from the contralateral paw in which MSU was injected. Light colors correspond to PBS paws, dark colors to MSU-injected paws. Symbols represent individual mice, horizontal lines and bars correspond to mean +/- SEM; Results were analyzed with a two-tailed Mann-Whitney test, * = p<0.05; ** = p < 0.01; *** = p < 0.001. In graphs A. and B. the area under curve (AUC) was determined and analyzed by a Mann-Whitney test.

## Discussion

### Inflammasomes triggered by MSU crystals and the nature of the priming inducer (signal I) *in vivo*

Gout is a frequent arthropathy with its etiology, i.e. MSU crystal deposition in the synovial fluid of patients, known for a long time (29). However, despite the discovery of NLRP3 as a Pattern Recognition Receptor sensing homeostatic disturbances and subsequent IL-1β secretion, the molecular mechanisms driving inflammatory manifestations upon MSU exposure are still poorly described. Several animal models have been developed to better understand the relationships between MSU crystals and IL-1β secretion, among which *ex vivo* cultures of macrophages were instrumental in the identification of the NLRP3 inflammasome. In addition, *in vivo* injections of crystals in the peritoneal cavity (8), into an artificial air pouch (30, 31) or directly inside the joint (32) have also been used to investigate neutrophil recruitment and inflammasome-dependent cytokine release. Most of these models exhibit important flaws and do not reproduce the complete pathophysiological manifestations of human gout or the joint microenvironment. For instance, inflammation triggered by intra-articular injection of MSU crystals in mice can be prevented by anti-TNFα therapy (33), which is inefficacious in humans. Here, we thoroughly characterized the impact of subcutaneous injection of MSU crystals in mouse paws, an approach (previously used by others (11, 13) that induces clinical manifestations (soft tissue inflammation with tendinitis, as seen with relevant medical imaging techniques) that can be preventable with drugs (colchicine, anakinra) currently in use to treat gout patients. Using this model in control and genetically-deficient mice, we demonstrated that, as opposed to previous observations in isolated macrophages, MSU-induced inflammation *in vivo* can occur independently from NLRP3, while being strictly IL-1β-dependent, suggesting that MSU- and silica-induced inflammation may proceed by different mechanisms (34). Therefore, we questioned whether other inflammasomes might also be involved in this process and possibly mask the NLRP3 effect. One current hypothesis postulates that neutrophils, following DNA release through NETosis (a process that can be activated by MSU crystals (23)), could provide a physiologically-relevant signal driving IL-1β secretion and inflammation. In this work, we tested the involvement of the AIM2 inflammasome and showed that mutant mice devoid of this DNA sensor, as well as animals in which both NLRP3 and AIM2 are lacking, develop almost normal local and systemic inflammatory responses following MSU crystals injection. Similar observations were made in *Casp 1*^*-/-*^; *11*^*-/-*^ animals. These experiments reveal the complexity of the MSU-induced inflammation and IL-1β secretion, which, in the absence of NLRP3 and AIM2, might likely be maturated by neutrophil-derived proteases (35). Furthermore, we also addressed the question of the nature of signal I, which is necessary to activate the NF-κB pathway and induce *Il-1β* transcription. Using co-cultures of neutrophils and macrophages, we showed that NETotic neutrophils cannot drive the secretion of IL-1β by MSU-stimulated macrophages, indicating that extracellular DNA is not capable to substitute for LPS in this process. Therefore, even though we were unable to identify a “physiologically”-relevant primer, our RNAseq analysis, which revealed *Tlr1, 2* and *6* overexpression upon MSU challenge, combined to experiments comparing *MyD88* and *Il-1R1* KO mice provide solid arguments supporting a role for endogenous lipids or lipopeptides (capable of triggering TLR1/2/6 signaling) as pathophysiological inducers of signal I-dependent NF-κB activation. Future work will aim at identifying such ligands.

### Signaling pathways supporting anti-inflammatory effects of type I IFNs

The anti-inflammatory effects of type I interferons are a matter of controversy, but can be supported by at least two lines of evidence: (i) sustained type I IFN production in chronic viral infections induces immunosuppression (36), as demonstrated in the LCMV model (37, 38). (ii) IFNβ is an effective treatment against inflammation of the central nervous system in multiple sclerosis (39). However, precise mechanisms underlying these immunomodulatory effects are still poorly understood. Using an alum-induced peritonitis model, the induction of the IL-10/STAT3 axis in response to type I IFN and LPS signaling was proposed as a predominant event to explain subsequent immunosuppression (26). In our dataset of imiquimod-modulated genes, several STAT3 targets (e.g. *Ccl2, Cxcr2, Socs3*) appeared significantly repressed, which indicates that this mechanism can be at play in response to imiquimod application. However, it is important to consider that our RNAseq approach does not enable the identification of protein modifications that could reveal the potential roles of transcription factors. A combined proteomic approach would help and probably refine the likely IL-10-dependent signaling and its contribution to imiquimod-based therapy. Furthermore, the timing (24 hpi) at which gene expression was quantified may also explain why transcripts encoding anti-inflammatory players, like IL-10, were not identified. Interestingly, *RUNX3*, another transcription factor involved in *Il-1β* gene regulation (27), was also evidenced in the analysis of our RNAseq data by the Reactome algorithm. Although this information requires further experimental validation, it is also supported by the mining of the ENCODE database which indicates that, in addition to *Il-1b*, multiple cytokines genes (such as *Il-6*) can be targeted by this transcription factor.

### Imiquimod, a powerful and promising alternative in gout patients

The present work clearly demonstrates the potential benefits to use imiquimod to treat gouty arthritis. Indeed, for several patients exhibiting additional comorbidities, such as kidney dysfunction, the prescription of colchicine might be unadvisable and imiquimod might become an attractive alternative devoid of major side effects. As demonstrated in our animal model, a single topical application was sufficient to reduce both neutrophils attraction and IL-1β secretion, thereby limiting the biological and clinical inflammatory symptoms that appear following MSU crystals injection. This dual activity of imiquimod appears even more attractive in view of the recent description of an inflammatory disorder (NOMID, neonatal-onset multisystem inflammatory disease) caused by GASDERMIN D (GSDMD) release and pyroptosis following inflammasome activation (40). Indeed, some patients suffering cryopyrin-associated periodic syndromes (CAPS) such as NOMID appear resistant to IL-1β blocking agents. A clinical trial will of course be necessary to evaluate the efficacy of imiquimod in humans. In addition, our work pointed to the multiplicity of actors (sensors, proteases) that are likely involved in the development of gout, which may advocate against the pharmacological targeting of only one of them, i.e the NLRP3 sensor. Finally, even though our genome-wide transcriptional analysis did not precisely identify the exact molecular pathways linking imiquimod, type I interferons and reduced *Il-1*β expression, an essential genetic network that appeared targeted by imiquimod (and predicted by IPA) brings together many genes encoding granulocyte cell surface markers (such as integrins), connected to a hub composed of the kinases SYK and PI3K. Their activation by MSU crystals binding to the plasma membrane (41) thus appears as an important contributor to neutrophil recruitment and activation, which could be impaired by already existing inhibitors developed for other pathologies, such as leukemia. Furthermore, RNAseq analysis using the ImmQuant software predicts that skin dendritic cells might be important mediators of the imiquimod-dependent anti-inflammatory response. The functional connection linking these cells and reduced neutrophil infiltrate will be a major research focus of ours.

In conclusion, our *in vivo* model uncovers an unsuspected complexity of the pathogenesis of gout triggered by urate crystals, as illustrated by its independence from the NLRP3 inflammasome. In addition, novel insights emerge from our transcriptomic analyses, such as the possible role of TLR1, 2 and/or 6 as sensors priming the NK-κB pathway. This observation paves the way for the search of lipids or lipoproteins which can be of endogenous or microbial (42) origin. Finally, our transcriptomic analyses reveal novel therapeutic targets, which will be necessary to develop in order to take care of an ever-growing number of patients in ageing populations.

## Materials and Methods

### Mice

*Nlrp3*^*-/-*^ and *Casp1/11*^*-/-*^ mice were provided by Romeo Ricci (Institut de Génétique et de Biologie Moléculaire (IGBMC), Illkirch, France), *Aim2*^*-/-*^ mice were obtained from Veit Hornung (Ludwig-Maximilians-Universität München, Germany) and *Ifnar1*^*-/-*^ from Rolf Zinkernagel (University Hospital, Zurich, Switzerland); we generated the double knock out mice (*Aim2*^*-/-*^; *Nlrp3*^*-/-*^) in our animal facility. *Il-1α* KO, *Il-1β* KO, *Il-1R1* KO and *MyD88* KO mice were provided by Bernhard Ryffel (UMR7355 CNRS - Université d’Orléans, France). Littermate controls are indicated by +/+ in all figures. All the mice used in these experiments were maintained under pathogen-free conditions in the animal facility of our laboratory (Institut d’Hématologie et d’Immunologie, Strasbourg, France) and were sex- and age-matched (12-16 weeks).

### Study approval

Handling of mice and experimental procedures were conducted in accordance with the French law for the protection of laboratory animals. The procedures were approved by the “service vétérinaire de la Préfecture du Bas-Rhin” (Strasbourg, France) and by the Regional ethical committee for animal experimentation (CREMEAS) of the Strasbourg University, under the authorization number 2018083014133041.

### Reagents

MSU crystals were generated as described (43). Briefly, we dissolved 1,68g of uric acid (Sigma-Aldrich) into 500mL of PBS containing 0,01M of NaOH by heating at 70°C, pH was adjusted to 7,1-7,2. Then, the uric acid solution was left at room temperature until the crystals formed under mild agitation. Crystals were then washed in ethanol, dried, weighted and thermally treated (250°C for an hour) and finally sonicated to obtain crystals <50µm in length. They were aliquoted in sterile PBS and frozen at −20°C until use. Poly-(dA:dT) and Lipofectamine 3000 were purchased from Invivogen and Invitrogen, respectively; LPS (*Salmonella abortus equi*), ATP and Colchicine from Sigma-Aldrich; ELISA kits (IL-1β, IL-6, TNFα, MPO) from Biotechne. Thioglycolate was home-made from brewer’s thioglycolate (BD). The various antibodies used for Western blot were from Biotechne (anti-mouse IL-1β, AF401, and secondary HAF-109) or from Abcam (anti-tubulin) and Sigma-Aldrich (anti-vinculin). Anti-Gr1, Anti-CD11b, anti-CD14 and 7-AAD were obtained from eBioscience. CRT0066101 was provided from the lab of Romeo Ricci; Anakinra was generously provided by Ommar S. Omarjee and Alexandre Belot (Lyon University, France) and Etanercept by Christian Von Frenckel (Liege University, Belgium).

### Gouty arthritis model

In this model, 3mg of preformed MSU crystals (described above) resuspended in 70µL of sterile PBS were injected by the subcutaneous route in the left hind paws of animals; the right paw was injected with PBS as control. Hind paw thickness was measured with a caliper and arthritis visual scores were established on the basis of tarsal/ankle oedema and erythema presence (0 = no arthritis, 1 = slight swelling and/or erythema, 2 = moderate swelling/erythema, 3 = severe oedema/erythema and 4 = excessive oedema spanning all over the paw). Visual scores were made by two independent experimenters. Mice were also monitored for body temperature.

### Topical application of imiquimod

Mice were anesthetized by intramuscular injection of Ketamine/Xylazine. The imiquimod-containing cream (ALDARA) or vehicle control cream were applied on the hind paws from the ankle to the tip of digits with a pen embedded into a latex glove. 250mg of ALDARA containing 5% imiquimod were sufficient to treat 10-12 paws, corresponding approximately to 1-1,25mg of imiquimod/paw (12).

### MSU-induced peritonitis model

One milligram of preformed MSU crystals was injected intraperitoneally at h0 and a peritoneal lavage was conducted 6h later, under anesthesia, with cold PBS. Peritoneal fluid was subsequently analyzed by flow cytometry and ELISA. Briefly, the liquid was filtered through a 40µm porosity strainer, centrifuged at 300g 5min at 4°C and cells were stained with anti-Gr1-FITC, anti-CD11b-APC and anti-CD14-PE antibodies to count the neutrophil and monocyte populations. Absolute numbers were also determined in peritoneal collections by multiplying the percentages of neutrophils/monocytes to the absolute numbers of cells in the lavage. Flow cytometry acquisitions were realized with the BD Accuri C6 cytometer (BD), data were analyzed with the BD Accuri Software.

### Magnetic resonance imagery (MRI)

To detect and localize inflammation in paws, anatomical MRI was acquired in eight ex-vivo mice. The day before MRI exam, 3mg of MSU and PBS were injected subcutaneously in the left and right paws respectively. ALDARA cream was applied on paws in four mice; control cream was applied on the other four. Mice were sacrificed under isoflurane anesthesia just before the MRI. MRI was performed on 7/30 Biospec system (Bruker Biospin, Ettlingen, Germany). Transmission was achieved with a quadrature volume resonator (inner diameter of 86 mm), and a surface coil (∼ 10 mm), installed on paw, was used for signal reception (Bruker BioSpin, Ettlingen, Germany). MRI experiments were realized with ParaVision 6.0.1 software. T2WI anatomical dataset was acquired using RARE-3D sequence using following parameters: FOV: 15 x 15 x 8 mm3, matrix 150 x 150 x 80, TE eff = 26.7 ms, TR = 2 s, N avg = 1, RARE-Factor = 14.

### MRI Data analysis and 3D reconstruction

The signal bias of T2WI induced by the surface coil used was corrected with N4 bias correction (Advanced Normalization Tools, ANTs). Paws were automatically co-registered based on the bones to one of the control mouse using FLIRT (FMRIB Software Library, Oxford, UK). The inflammatory areas were automatically segmented using FAST (FMRIB Software Library, Oxford, UK). Volumetric images were obtained with the help of the 3D viewer module of Image J, 3D reconstruction was made using the 3D Slicer software.

### Protein preparation from paws

In order to quantify the cytokine production *in situ*, hind paws were collected from euthanized mice and stored at −80°C. Paws were first minced with a scalpel in a Petri dish placed on dry ice. Small tissue pieces were maintained on ice and 1mL of NP40 lysis buffer was added before homogenization with a tissue tearer (OMNI International Tissue MASTER 125). Tubes were then kept on ice for 45min and centrifuged at 12 000g 20min at 4°C; afterwards, the supernatants were collected in individual Eppendorf protein-low bind 1,5mL tubes (Eppendorf) and stored at −80°C. Cytokine measurements were performed by ELISA and Western Blot.

### RNA preparation from paws

RNAs were prepared similarly to proteins (collection of paws, slicing) but tissues were homogenized in 2mL of Trizol (TRI-Reagent, Sigma-Aldrich) instead of NP40 lysis buffer. Homogenates were transferred in a nuclease-free 1,5mL Eppendorf tube and RNA was purified as previously described (12).Total RNAs were quantified using a Nanodrop device and then treated with DNAse I (Roche) according to the manufacturer’s instructions.

### Real-time quantitative PCR (RT-qPCR)

Total RNA was reverse transcribed using the cDNA synthesis kit (iScript ready-to-use cDNA supermix, Biorad). Real-time quantitative RT-qPCR was performed in a total volume of 20µL using the Sso-advanced universal SYBR-Green supermix (Biorad) and gene-specific primers (sequence available upon request). After a denaturing step at 95°C for 30 seconds, 40 cycles were performed (95°C for 5s and 60°C for 20s) using a Rotor-Gene 6000 real-time PCR machine (Corbett Life Science). Results were obtained using the SDS Software (Perkin Elmer) and evaluated using Excel (Microsoft). Melting-curve analysis was performed to assess the specificity of PCR products. Relative expression was calculated using the comparative threshold cycle (Ct) method, normalized on a control group.

### RNA-seq

Total RNA integrity was determined with the Agilent total RNA Pico Kit on a 2100 Bioanalyzer instrument (Agilent Technologies, Paolo Alto, USA). Library construction was performed with the “SMARTer^®^ Stranded Total RNA-Seq Kit v2 - Pico Input Mammalian” (TaKaRa Bio USA, Inc., Mountain View, CA, USA) with a final multiplexing of 11 libraries according to the manufacturer’s instructions. The library pool was denatured according to the Illumina protocol “Denature and Dilute Libraries Guide” and then deposited at a final concentration of 1.3 pM to be sequenced on the NextSeq 500 (Illumina Inc., San Diego, CA, USA).

### Analysis of RNA-seq data

The transcriptome data set, composed of sequencing reads, was generated by an Illumina NextSeq instrument. The objective was to identify genes that are differentially expressed between two experimental conditions, namely treatment and control. For every sample, quality control was carried out and assessed with the NGS Core Tools FastQC (http://www.bioinformatics.babraham.ac.uk/projects/fastqc/). Sequence reads are mapped using STAR (44) and unmapped reads are remapped with Bowtie2 (45) using with the very sensitive local option. The total reads mapped were finally available in BAM (Binary Alignment Map) format for raw read counts extraction. Read counts were found by the htseq-count tool of the Python package HTSeq (46) with default parameters to generate an abundant matrix. At last, differential analyses were performed by the DESEQ2 (47) package of the Bioconductor framework. Up-regulated and down-regulated genes were selected based on the adjusted p-value and the fold-change information. Gene expression was analyzed using dedicated R scripts to build volcano plots and graphs of Gene Ontology (GO) terms enrichment. Heatmaps were built using the online application heatmapper (http://www.heatmapper.ca/). IPA (Ingenuity pathway analysis, Qiagen) was used for pathway analysis. Predictions of immune cell population from RNAseq data were made with the ImmQuant deconvolution software (22).

### Western blot

IL-1β-maturation quantification was performed by western blot from ground paw samples (see previous sections). Briefly, total proteins were quantified in paw extracts using the Bicinchoninic acid technique (BCA) and 75-100µg of proteins were loaded on 12% polyacrylamide gels (Biorad). The migration step was followed by a liquid transfer on PVDF membranes (100V 40min) and a blocking step for 1h with TBS-Tween 1% milk 5%. Membranes were then probed with AF-401 (IL-1β), anti-vinculine (V9131, Sigma-Aldrich) or anti-tubulin (DM1A, Abcam) antibodies at 4°C overnight. HAF-109 and goat anti-mouse antibodies were subsequently used as secondary antibodies (1h, RT). Revelation was conducted with the Femto super-signal kit. Three washes were realized between each stage.

### Cell culture

Peritoneal macrophages were collected from peritoneal exudates 72h after the injection of 2mL of Thioglycolate 4%. Cells were washed, counted and plated at a density of 5×10^5^ cells/well (24 well-plates) in RPMI 1640 (Gibco) 10% FBS (Dutcher); the medium was changed for a fresh one 3h later. Peritoneal neutrophils were harvested after a short course of thioglycolate (18h) and purified using Ficoll Plaque leading to a purity greater than 90%. Neutrophils were plated immediately at a density of 1×10^6^ cells/well in RPMI 1640 10% FCS.

### Inflammasome activation

In order to stimulate the production of pro-IL-1β, cells were pre-treated with LPS 1µg/mL for 3h and the NLRP3 inflammasome was then activated with 2mM ATP for 1h or with MSU crystals (250µg/mL) for 6h. The AIM2 inflammasome was activated by the transfection of 0.5µg of poly-dA:dT using the Lipofectamine 3000 kit, the medium was collected 8h after transfection. After each activation, the supernatants were collected, centrifuged at 2000g 5min 4°C to get rid of the cell debris and the supernatants were harvested in new tubes, all samples were kept a −80°C.

### Inhibitors

The CRT0066101 inhibitor was used *in vitro* at a dose of 10µM and introduced in the culture medium 1h prior to LPS priming and kept at this concentration all along the activation. 0.1 % DMSO was used as control.

### Statistics

Data were analyzed with a Mann-Whitney test (two-tailed unpaired) to compare two independent groups using GraphPad 5.01 software. A probability (p) value of < 0.05 was considered significant. *p < 0.05, **p < 0.01, ***p < 0.001.

## Author contributions

AM conducted, acquired and analyzed data and wrote the manuscript; ADC acquired and analyzed data; CP acquired and analyzed data; CAF, AP and CM acquired data; NP, IA, RC, EC, JS and JPA analyzed data; BF provided reagents; SB analyzed data and wrote the manuscript; PG designed research studies, analyzed data and wrote the manuscript

## Acknowledgements

Work in our laboratory is supported by grants from the Agence Nationale de la Recherche (ANR) (ANR-11-LABX-0070_TRANSPLANTEX), the INSERM (UMR_S 1109), the Institut Universitaire de France (IUF), the University of Strasbourg (IDEX UNISTRA), the European regional development fund (European Union) INTERREG V program (project n°3.2 TRIDIAG) and MSD-Avenir grant AUTOGEN. The input and advices from Roméo Ricci and Zhirong Zhang (IGBMC, Illkirch, France) at the initial step of this work is greatly acknowledged. PG thanks Dr. Stephan Blüml (Vienna Medical University, Austria) for helpful discussions and Dr. Philippe Bouillet (WEHI, Melbourne, Australia) for critical reading of the manuscript.

## References

1. Elfishawi, M.M., Zleik, N., Kvrgic, Z., Michet, C.J., Jr., Crowson, C.S., Matteson, E.L., and Bongartz, T. 2017. The Rising Incidence of Gout and the Increasing Burden of Comorbidities: A Population-based Study over 20 Years. J Rheumatol.

2. Nakayama, A., Nakaoka, H., Yamamoto, K., Sakiyama, M., Shaukat, A., Toyoda, Y., Okada, Y., Kamatani, Y., Nakamura, T., Takada, T., et al. 2017. GWAS of clinically defined gout and subtypes identifies multiple susceptibility loci that include urate transporter genes. Ann Rheum Dis 76:869-877.

3. Dalbeth, N., Stamp, L.K., and Merriman, T.R. 2017. The genetics of gout: towards personalised medicine? BMC Med 15:108.

4. Chen, C.J., Tseng, C.C., Yen, J.H., Chang, J.G., Chou, W.C., Chu, H.W., Chang, S.J., and Liao, W.T. 2018. ABCG2 contributes to the development of gout and hyperuricemia in a genomewide association study. Sci Rep 8:3137.

5. Kottgen, A., Albrecht, E., Teumer, A., Vitart, V., Krumsiek, J., Hundertmark, C., Pistis, G., Ruggiero, D., O’Seaghdha, C.M., Haller, T., et al. 2013. Genome-wide association analyses identify 18 new loci associated with serum urate concentrations. Nat Genet 45:145-154.

6. Kuemmerle-Deschner, J.B., Ozen, S., Tyrrell, P.N., Kone-Paut, I., Goldbach-Mansky, R., Lachmann, H., Blank, N., Hoffman, H.M., Weissbarth-Riedel, E., Hugle, B., et al. 2017. Diagnostic criteria for cryopyrin-associated periodic syndrome (CAPS). Ann Rheum Dis 76:942-947.

7. So, A.K., and Martinon, F. 2017. Inflammation in gout: mechanisms and therapeutic targets. Nat Rev Rheumatol 13:639-647.

8. Martinon, F., Petrilli, V., Mayor, A., Tardivel, A., and Tschopp, J. 2006. Gout-associated uric acid crystals activate the NALP3 inflammasome. Nature 440:237-241.

9. Giamarellos-Bourboulis, E.J., Mouktaroudi, M., Bodar, E., van der Ven, J., Kullberg, B.J., Netea, M.G., and van der Meer, J.W. 2009. Crystals of monosodium urate monohydrate enhance lipopolysaccharide-induced release of interleukin 1 beta by mononuclear cells through a caspase 1-mediated process. Ann Rheum Dis 68:273-278.

10. Maueroder, C., Kienhofer, D., Hahn, J., Schauer, C., Manger, B., Schett, G., Herrmann, M., and Hoffmann, M.H. 2015. How neutrophil extracellular traps orchestrate the local immune response in gout. J Mol Med (Berl) 93:727-734.

11. Schauer, C., Janko, C., Munoz, L.E., Zhao, Y., Kienhofer, D., Frey, B., Lell, M., Manger, B., Rech, J., Naschberger, E., et al. 2014. Aggregated neutrophil extracellular traps limit inflammation by degrading cytokines and chemokines. Nat Med 20:511-517.

12. Nehmar, R., Alsaleh, G., Voisin, B., Flacher, V., Mariotte, A., Saferding, V., Puchner, A., Niederreiter, B., Vandamme, T., Schabbauer, G., et al. 2017. Therapeutic Modulation of Plasmacytoid Dendritic Cells in Experimental Arthritis. Arthritis Rheumatol 69:2124-2135.

13. Yang, G., Yeon, S.H., Lee, H.E., Kang, H.C., Cho, Y.Y., Lee, H.S., and Lee, J.Y. 2018. Suppression of NLRP3 inflammasome by oral treatment with sulforaphane alleviates acute gouty inflammation. Rheumatology (Oxford) 57:727-736.

14. Liebner, R., Mathaes, R., Meyer, M., Hey, T., Winter, G., and Besheer, A. 2014. Protein HESylation for half-life extension: synthesis, characterization and pharmacokinetics of HESylated anakinra. Eur J Pharm Biopharm 87:378-385.

15. Amaral, F.A., Costa, V.V., Tavares, L.D., Sachs, D., Coelho, F.M., Fagundes, C.T., Soriani, F.M., Silveira, T.N., Cunha, L.D., Zamboni, D.S., et al. 2012. NLRP3 inflammasome-mediated neutrophil recruitment and hypernociception depend on leukotriene B(4) in a murine model of gout. Arthritis Rheum 64:474-484.

16. Castelblanco, M., Lugrin, J., Ehirchiou, D., Nasi, S., Ishii, I., So, A., Martinon, F., and Busso, N. 2018. Hydrogen sulfide inhibits NLRP3 inflammasome activation and reduces cytokine production both in vitro and in a mouse model of inflammation. J Biol Chem 293:2546-2557.

17. Joosten, L.A., Ea, H.K., Netea, M.G., and Busso, N. 2011. Interleukin-1beta activation during acute joint inflammation: a limited role for the NLRP3 inflammasome in vivo. Joint Bone Spine 78:107-110.

18. Joosten, L.A., Netea, M.G., Mylona, E., Koenders, M.I., Malireddi, R.K., Oosting, M., Stienstra, R., van de Veerdonk, F.L., Stalenhoef, A.F., Giamarellos-Bourboulis, E.J., et al. 2010. Engagement of fatty acids with Toll-like receptor 2 drives interleukin-1beta production via the ASC/caspase 1 pathway in monosodium urate monohydrate crystal-induced gouty arthritis. Arthritis Rheum 62:3237-3248.

19. Zhang, Z., Meszaros, G., He, W.T., Xu, Y., de Fatima Magliarelli, H., Mailly, L., Mihlan, M., Liu, Y., Puig Gamez, M., Goginashvili, A., et al. 2017. Protein kinase D at the Golgi controls NLRP3 inflammasome activation. J Exp Med 214:2671-2693.

20. Chen, K.W., Bezbradica, J.S., Gross, C.J., Wall, A.A., Sweet, M.J., Stow, J.L., and Schroder, K. 2016. The murine neutrophil NLRP3 inflammasome is activated by soluble but not particulate or crystalline agonists. Eur J Immunol 46:1004-1010.

21. Soares, D.M., Figueiredo, M.J., Martins, J.M., Machado, R.R., Kanashiro, A., Malvar Ddo, C., Pessini, A.C., Roth, J., and Souza, G.E. 2009. CCL3/MIP-1 alpha is not involved in the LPS-induced fever and its pyrogenic activity depends on CRF. Brain Res 1269:54-60.

22. Frishberg, A., Brodt, A., Steuerman, Y., and Gat-Viks, I. 2016. ImmQuant: a user-friendly tool for inferring immune cell-type composition from gene-expression data. Bioinformatics 32:3842-3843.

23. Chatfield, S.M., Grebe, K., Whitehead, L.W., Rogers, K.L., Nebl, T., Murphy, J.M., and Wicks, I.P. 2018. Monosodium Urate Crystals Generate Nuclease-Resistant Neutrophil Extracellular Traps via a Distinct Molecular Pathway. J Immunol 200:1802-1816.

24. Lugrin, J., and Martinon, F. 2018. The AIM2 inflammasome: Sensor of pathogens and cellular perturbations. Immunol Rev 281:99-114.

25. Thacker, S.G., Berthier, C.C., Mattinzoli, D., Rastaldi, M.P., Kretzler, M., and Kaplan, M.J. 2010. The detrimental effects of IFN-alpha on vasculogenesis in lupus are mediated by repression of IL-1 pathways: potential role in atherogenesis and renal vascular rarefaction. J Immunol 185:4457-4469.

26. Guarda, G., Braun, M., Staehli, F., Tardivel, A., Mattmann, C., Forster, I., Farlik, M., Decker, T., Du Pasquier, R.A., Romero, P., et al. 2011. Type I interferon inhibits interleukin-1 production and inflammasome activation. Immunity 34:213-223.

27. Lim, B., Ju, H., Kim, M., and Kang, C. 2011. Increased genetic susceptibility to intestinal-type gastric cancer is associated with increased activity of the RUNX3 distal promoter. Cancer 117:5161-5171.

28. Deguine, J., and Barton, G.M. 2014. MyD88: a central player in innate immune signaling. F1000Prime Rep 6:97.

29. McCarty, D.J., and Hollander, J.L. 1961. Identification of urate crystals in gouty synovial fluid. Ann Intern Med 54:452-460.

30. Akahoshi, T., Namai, R., Murakami, Y., Watanabe, M., Matsui, T., Nishimura, A., Kitasato, H., Kameya, T., and Kondo, H. 2003. Rapid induction of peroxisome proliferator-activated receptor gamma expression in human monocytes by monosodium urate monohydrate crystals. Arthritis Rheum 48:231-239.

31. Ryckman, C., McColl, S.R., Vandal, K., de Medicis, R., Lussier, A., Poubelle, P.E., and Tessier, P.A. 2003. Role of S100A8 and S100A9 in neutrophil recruitment in response to monosodium urate monohydrate crystals in the air-pouch model of acute gouty arthritis. Arthritis Rheum 48:2310-2320.

32. Joosten, L.A., Crisan, T.O., Azam, T., Cleophas, M.C., Koenders, M.I., van de Veerdonk, F.L., Netea, M.G., Kim, S., and Dinarello, C.A. 2016. Alpha-1-anti-trypsin-Fc fusion protein ameliorates gouty arthritis by reducing release and extracellular processing of IL-1beta and by the induction of endogenous IL-1Ra. Ann Rheum Dis 75:1219-1227.

33. Amaral, F.A., Bastos, L.F., Oliveira, T.H., Dias, A.C., Oliveira, V.L., Tavares, L.D., Costa, V.V., Galvao, I., Soriani, F.M., Szymkowski, D.E., et al. 2016. Transmembrane TNF-alpha is sufficient for articular inflammation and hypernociception in a mouse model of gout. Eur J Immunol 46:204-211.

34. Benmerzoug, S., Rose, S., Bounab, B., Gosset, D., Duneau, L., Chenuet, P., Mollet, L., Le Bert, M., Lambers, C., Geleff, S., et al. 2018. STING-dependent sensing of self-DNA drives silica-induced lung inflammation. Nat Commun 9:5226.

35. Joosten, L.A., Netea, M.G., Fantuzzi, G., Koenders, M.I., Helsen, M.M., Sparrer, H., Pham, C.T., van der Meer, J.W., Dinarello, C.A., and van den Berg, W.B. 2009. Inflammatory arthritis in caspase 1 gene-deficient mice: contribution of proteinase 3 to caspase 1-independent production of bioactive interleukin-1beta. Arthritis Rheum 60:3651-3662.

36. Snell, L.M., McGaha, T.L., and Brooks, D.G. 2017. Type I Interferon in Chronic Virus Infection and Cancer. Trends Immunol 38:542-557.

37. Wang, Y., Swiecki, M., Cella, M., Alber, G., Schreiber, R.D., Gilfillan, S., and Colonna, M. 2012. Timing and magnitude of type I interferon responses by distinct sensors impact CD8 T cell exhaustion and chronic viral infection. Cell Host Microbe 11:631-642.

38. Teijaro, J.R., Ng, C., Lee, A.M., Sullivan, B.M., Sheehan, K.C., Welch, M., Schreiber, R.D., de la Torre, J.C., and Oldstone, M.B. 2013. Persistent LCMV infection is controlled by blockade of type I interferon signaling. Science 340:207-211.

39. Reich, D.S., Lucchinetti, C.F., and Calabresi, P.A. 2018. Multiple Sclerosis. N Engl J Med 378:169-180.

40. Xiao, J., Wang, C., Yao, J.C., Alippe, Y., Xu, C., Kress, D., Civitelli, R., Abu-Amer, Y., Kanneganti, T.D., Link, D.C., et al. 2018. Gasdermin D mediates the pathogenesis of neonatal-onset multisystem inflammatory disease in mice. PLoS Biol 16:e3000047.

41. Ng, G., Sharma, K., Ward, S.M., Desrosiers, M.D., Stephens, L.A., Schoel, W.M., Li, T., Lowell, C.A., Ling, C.C., Amrein, M.W., et al. 2008. Receptor-independent, direct membrane binding leads to cell-surface lipid sorting and Syk kinase activation in dendritic cells. Immunity 29:807-818.

42. Liu, J., Cui, L., Yan, X., Zhao, X., Cheng, J., Zhou, L., Gao, J., Cao, Z., Ye, X., and Hu, S. 2018. Analysis of Oral Microbiota Revealed High Abundance of Prevotella Intermedia in Gout Patients. Cell Physiol Biochem 49:1804-1812.

43. Schiltz, C., Liote, F., Prudhommeaux, F., Meunier, A., Champy, R., Callebert, J., and Bardin, T. 2002. Monosodium urate monohydrate crystal-induced inflammation in vivo: quantitative histomorphometric analysis of cellular events. Arthritis Rheum 46:1643-1650.

44. Dobin, A., Davis, C.A., Schlesinger, F., Drenkow, J., Zaleski, C., Jha, S., Batut, P., Chaisson, M., and Gingeras, T.R. 2013. STAR: ultrafast universal RNA-seq aligner. Bioinformatics 29:15-21.

45. Langmead, B., and Salzberg, S.L. 2012. Fast gapped-read alignment with Bowtie 2. Nat Methods 9:357-359.

46. Anders, S., Pyl, P.T., and Huber, W. 2015. HTSeq--a Python framework to work with high-throughput sequencing data. Bioinformatics 31:166-169.

47. Love, M.I., Huber, W., and Anders, S. 2014. Moderated estimation of fold change and dispersion for RNA-seq data with DESeq2. Genome Biol 15:550.

