## Supplementary material for "NLRP3- and AIM2-autonomy in a mouse model of MSU crystal-induced acute inflammation *in vivo* highlights imiquimod-dependent targeting of *Il-1β* expression as relevant therapy for gout patients": supl Fig S1 to S14

**Suppl. Fig. S1. The murine subcutaneous acute uratic inflammation model is clinically similar to human gout. A and B.** Representative pictures of *C57BL/6J* mice paws taken 24h (acute phase) and 168h (Resolution phase) after the injection of 3mg of MSU crystals. Paws are typically inflamed at the early phase (R: redness; Oe: Oedema; AOe: Ankle oedema; T: Tophus/i); cardinal signs of inflammation tends to disappear at the resolution phase enabling numerous tophus-like structures (T) to be visible. **C.** Magnetic Resonance Imaging (MRI) Axial view of PBS (left) and MSU-injected (right) paws of *C57BL/6J* mice at 24h post-injection. MRI was performed with the T2-weighted, fat-suppression mode and reveals oedema and tenosynovitis upon MSU injection. **D.** MSU injection induces mature IL-1β production. Mice paws were collected at different time points after the injection of MSU crystals or PBS. Paw extracts were prepared and analyzed by western blot for IL-1β; Vinculin was used as loading control. The picture is representative of three different blots. * denotes non-specific bands.

**Suppl. Fig. S2. Subcutaneous acute uratic inflammation responds to the first-line drug regiment used in human gout. A-D.** *C57BL/6J* mice were submitted to an acute uratic inflammation experiment and treated with colchicine at day 0 (1mg/kg/j) and at day 1 (0,5mg/kg/j). **A.** Paw swelling measurements. **B.** Clinical scores. **C.** representative pictures of the colchicine-treated (right) and vehicle-treated (left) mice paws and **D.** Body temperature. **E-H.** Acute uratic inflammation (*C57BL/6J* mice) following Anakinra (100mg/kg/j) at D0 and D1 (**E-F**) or Etanercept treatment (10mg/kg) at D0 (**G-H**). **E.** and **G.** Paw swelling, **F.** and **H.** Clinical scores. All experiments were realized with n = 5 mice in each group. Results represent mean +/- SEM and were analyzed with a two-tailed Mann-Whitney test, * = $p<0.05$, ** = $p < 0.01$.

**Suppl. Fig. S3. Subcutaneous acute uratic inflammation is IL-1β-dependent.** Controls (*C57Bl/6*, green dots n=7) and $Il\text{-}1\beta^{-/-}$ (red dots n=9) and $Il\text{-}1\alpha^{-/-}$ (orange dots n=6) mice were injected s.c with MSU crystals and clinical signs were recorded during 24h. **A.** Paw swelling measurements. **B.** Clinical scores. At 24h, paws were dissected and proteins were analyzed by ELISA. **C.** IL-1β, **D.** IL-6 and **E.** MPO quantification. Symbols represent individual mice with light colored dots representing PBS paws and bright colored dots the MSU-injected paws. Horizontal lines and bars correspond to mean +/- SEM; Results were analyzed with a two-tailed Mann-Whitney test, * = $p<0.05$, ** = $p < 0.01$, *** = $p < 0.001$, ns = not significant. In graphs A. and B. the area under curve (AUC) was determined and analyzed by a Mann-Whitney test.

**Suppl. Fig. S4. Pharmacological inhibition of the PKD-NLRP3 axis phenocopies the *Nlrp3* knock out. C57BL/6J** mice were treated with the Pan-PKD inhibitor CRT0066101 (10mg/kg; n=5) or the corresponding vehicle (DMSO; n=4) prior the induction of acute uratic inflammation. CRT0066101 or vehicle were then injected every 12h (10mg/kg). **A-C.** Clinical data were recorded over 2 days. **A.** Paw swelling, **B.** clinical scores and **C.** Body temperature. **D.-E.** Paw extracts were analyzed by western blot to reveal IL-1β (**D.**) and ELISA quantifications of total IL-1β (**E.**). **A-C**. data are representative of 3 independent experiments. **D-E**. n = 4 in each group. Symbols represent individual mice, horizontal lines and bars correspond to mean +/- SEM; Results were analyzed with a two-tailed Mann-Whitney test, ns = not significant.

**Suppl. Fig. S5. IL-1β maturation depends on the PKD-NLRP3 axis *in vitro*. A.** Peritoneal macrophages isolated from $Nlrp3^{+/+}$ (n=4) and $Nlrp3^{-/-}$ (n=4) mice were seeded at $5 \times 10^5$ cells/well and stimulated with lipopolysaccharide only (LPS, 1µg/mL) or LPS plus MSU or ATP. $Nlrp3^{+/+}$ macrophages were treated with the CRT0066101 inhibitor (yellow dots) or DMSO (green dots) 1h prior and all along the stimulation; DMSO-treated *Nlrp3* $^{-/-}$ macrophages were used as controls (red dots). IL-1β was quantified by ELISA. **B.** The same experiment as in **A.** was realized with $1x10^6$ murine neutrophils/well. **C.** Peritoneal macrophages from *Nlrp3*$^{+/+}$ (green dots) or *Nlrp3*$^{-/-}$ (red dots) were stimulated with LPS only or LPS plus MSU or ATP. TNFα and IL-6 were quantified by ELISA. The data are representative of 3 experiments. Symbols represent individual mice, horizontal lines and bars correspond to mean +/- SEM; Results were analyzed with a two-tailed Mann-Whitney test, * = p<0.05, ns = not significant.

**Suppl. Fig. S6. NLRP3 is not required in the MSU-induced peritonitis model.** *Nlrp3*$^{+/+}$ (n=8) and Nlrp3$^{-/-}$ mice (n=6) were subjected to a peritonitis model by injection of MSU crystals (1 mg) into the peritoneal cavity. **A-D.** Flow cytometry analysis of peritoneal exudate. **A.** Gating strategy: peritoneal cells were distinguished from debris according to FSC/SSC profile and live cells were selected. CD14+Gr1- were assigned to monocytes; neutrophils were defined as $Gr1^+CD11b^+$. **B.** Total cells according to the FSC/SSC profile, **C.** Percentage and absolute number of neutrophils, **D.** Percentage and absolute number of monocytes. **E.** ELISA measurements of peritoneal IL-1β (4 animals per group). *Nlrp3*$^{+/+}$ mice are depicted by green dots, *Nlrp3*$^{-/-}$ by red dots. Symbols represent individual mice, horizontal lines and bars correspond to mean +/- SEM; Results were analyzed with a two-tailed Mann-Whitney test, ns = not significant.

**Suppl. Fig. S7. RNAseq analysis of PBS *vs* MSU-injected paws.** 24h post injection, paws were harvested, RNA extracted and analyzed by RNAseq. **A.** Volcano plot with green dots showing significantly (p<0.05) upregulated genes (log2FC>2) and red dots significantly downregulated genes (log2FC<2). **B.** Heatmap showing unsupervised clusterization of the 405 most differentially modulated genes (p<0.005). **C.** Heatmap of the top 100 genes most significantly modulated by MSU injection. Genes discussed in the text are framed in red. **D.** Gene Ontology (GO) terms enrichment analysis.

**Suppl. Fig. S8. Neutrophil-associated genes are upregulated by MSU crystals and downregulated by topical imiquimod treatment.** An acute uratic inflammation experiment was conducted on two cohorts of animals (one paw was injected with MSU crystals and the contralateral one with PBS); one cohort was treated with topical imiquimod (Aldara cream), the other with a control cream. 24h after, paws were collected and total RNA were extracted and submitted to a genome-wide transcriptomic analysis by RNAseq. The most differentially expressed genes (p<0.001) in the two settings (**A.** MSU crystal-injected *vs* PBS-injected paws, **B.** MSU-injected and imiquimod-treated *vs* MSU-injected and control cream-treated paws) were analyzed with IPA (Ingenuity pathway analysis, Qiagen) software. Genes associated with neutrophil functions were significantly overexpressed following MSU injection and downregulated when topical imiquimod was applied. Solid bars/arrows indicate direct interactions, broken bars/arrows the indirect ones; green pictograms represent down-modulated genes, red ones overexpressed genes. For more information, see http://qiagen.force.com/KnowledgeBase/articles/Basic_Technical_Q_A/Legend.

**Suppl. Fig. S9. Neutrophil infiltrate induced by MSU injection is reduced by imiquimod topical application.** RNAseq data (normalized and log2 transformed) were analyzed by the ImmQuant software (http://csgi.tau.ac.il/ImmQuant/) which predicts 203 immune cell types enrichment in complex environments. **A.** Comparison of MSU- *vs* PBS-injected paws. **B.** Comparison of imiquimod-

*vs* control cream-treated paws after PBS injection. **C.** Comparison of imiquimod- *vs* control cream-treated paws after MSU injection. Cell enrichment is shown in red, exhaustion in blue. Only immune cells (dendritic cells DC, monocytes MO, macrophages MF and granulocytes GN) are shown in the figure. Some specific cell types discussed in the text are framed in red. **D.** Quantification of the data shown in **A-C** performed by ImmQuant. * = FDR<0.05.

**Suppl. Fig. S10. Caspases 1/11 have a limited impact in the subcutaneous MSU-induced acute uratic inflammation model.** *Casp1*$^{-/-}$ (bearing a passenger mutation compromising *Casp11* function) were submitted to an acute uratic inflammation experiment (n = 3 to 7) and clinical observations were made over a 48h period. **A.** Paw thickness, **B.** Clinical score, **C.** Body temperature and **D.** seric IL-1β measured ELISA. Symbols represent individual mice with green dots representing *Casp1*$^{+/+}$ mice and blue dots *Casp1*$^{-/-}$ (*Casp1*$^{-/-}$;*Casp11*$^{-/-}$) mice. Horizontal lines and bars correspond to mean +/- SEM; Results were analyzed with a two-tailed Mann-Whitney test, * = $p<0.05$, ns = not significant. In graph B, the area under curve (AUC) was determined and analyzed with a Mann-Whitney test.

**Suppl. Fig. S11. Topical imiquimod reduces subcutaneous oedemas and tenosynovitis.** MRI sequences performed as described in Figure 5 (MRI at 24hpi, T2, fat suppression) are presented (4 mice per group). Left, control cream-treated paws (5 pictures for 4 mice), the two upper pictures are from the same mouse and depict the concomitant presence of large subcutaneous oedemas and a marked tenosynovitis of the flexor *tendon communis* of the fingers; right, topical imiquimod-treated paws (4 pictures for 4 mice). Anatomic details are indicated by arrows (see legend in the lower-right corner). Contrast has been set manually for each pair of paws (PBS and MSU for each mouse). Each MSU paw should then be considered with regard to the PBS paw.

**Suppl. Fig. S12. RNAseq analysis of control (ctr) *vs* imiquimod-treated PBS-injected paws.** 24h post injection, paws were harvested, RNA extracted and analyzed by RNAseq. **A.** Volcano plot with green dots showing significantly ($p<0.05$) upregulated genes (log2FC>2) and red dots significantly downregulated genes (log2FC<2). **B.** Heatmap showing unsupervised clusterization of the 395 most differentially modulated genes ($p<0.005$). **C.** Heatmap of the top 100 genes most significantly modulated by imiquimod treatment upon PBS injection. Genes discussed in the text are framed in red. **D.** Gene Ontology (GO) terms enrichment analysis.

**Suppl. Fig. S13. RNAseq analysis of control (ctr) *vs* imiquimod-treated MSU-injected paws.** 24h post injection, paws were harvested, RNA extracted and analyzed by RNAseq. **A.** Volcano plot with green dots showing significantly ($p<0.05$) upregulated genes (log2FC>2) and red dots significantly downregulated genes (log2FC<2). **B.** Heatmap showing unsupervised clusterization of the 1118 most differentially modulated genes ($p<0.005$). **C.** Heatmap of the top 100 genes most significantly modulated by imiquimod treatment upon MSU injection. Genes discussed in the text are framed in red. **D.** Gene Ontology (GO) terms enrichment analysis.

**Suppl. Fig. S14. Topical imiquimod application affects a constellation of signaling pathway related to neutrophil adhesion and Syk-PI3K axis.** The 666 most differentially expressed genes between MSU-control cream- and MSU-imiquimod-treated paws found by RNAseq (the same data set used in Suppl. Fig. S8) were analyzed with IPA (Ingenuity pathway analysis, Qiagen) software. The main signaling pathways affected by topical imiquimod are depicted and connexions are shown in this gene map. Solid bars/arrows indicate direct interactions, broken bars/arrows the indirect ones; green pictograms represent down-modulated genes, red ones the overexpressed genes in imiquimod-treated relative to control-cream treated paws. For more information, see http://qiagen.force.com/KnowledgeBase/articles/Basic_Technical_Q_A/Legend.

Supplm Fig. S1

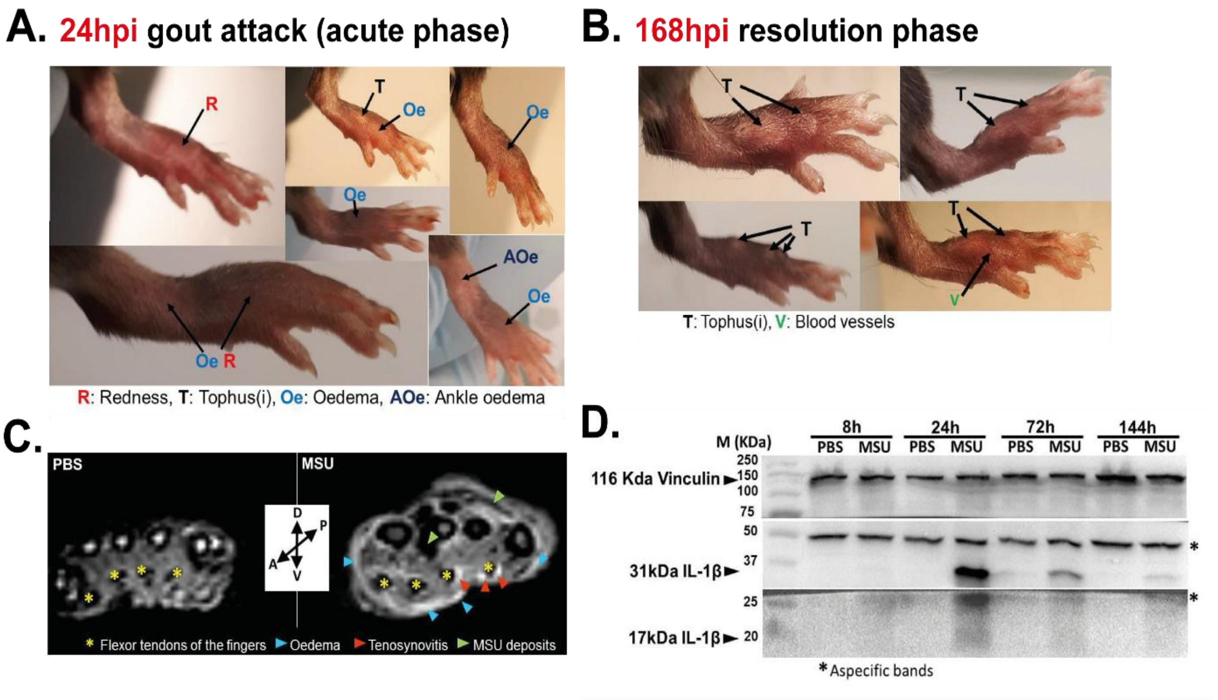

Supplm. Fig. S2

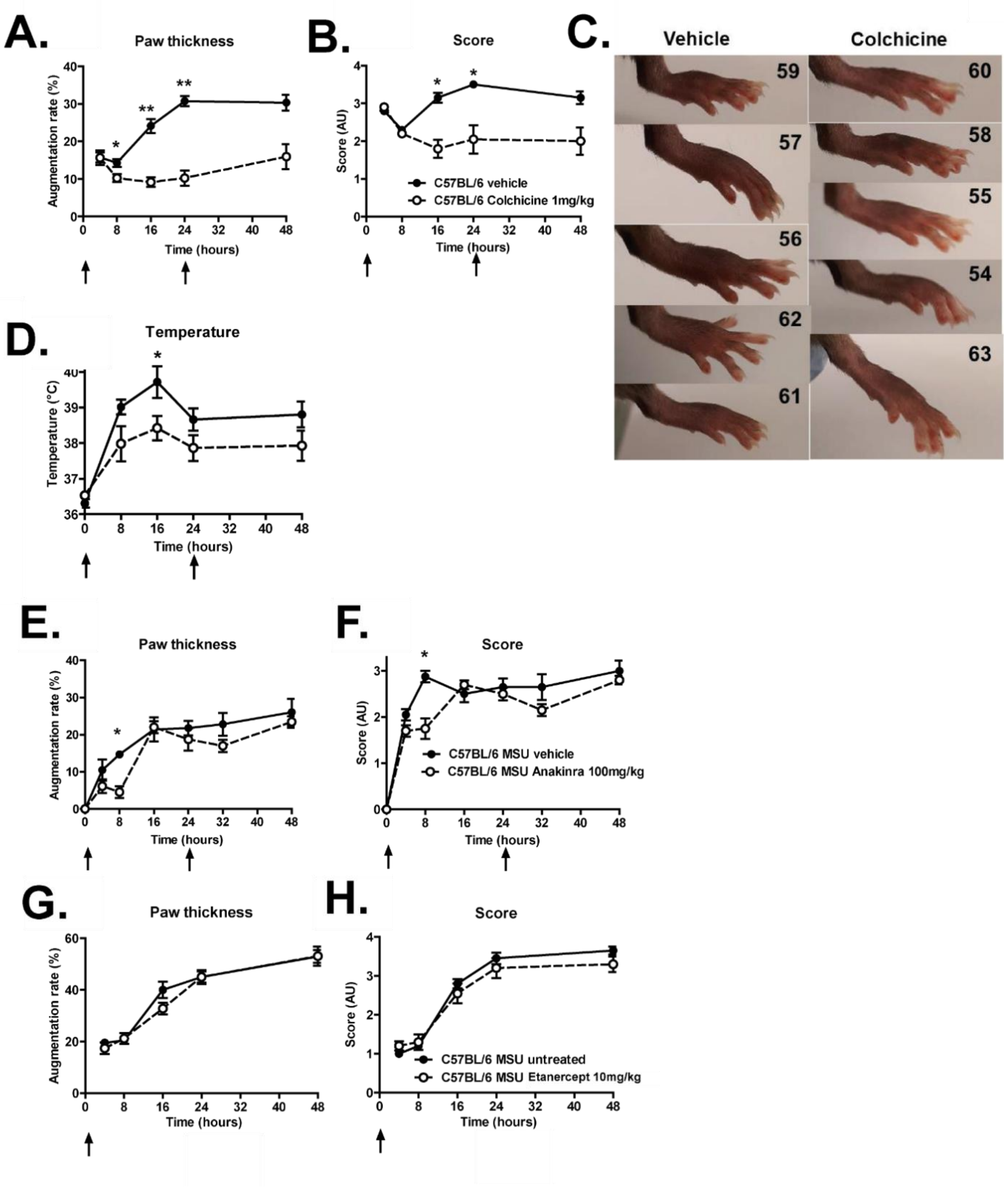

Supplm. Fig. S3

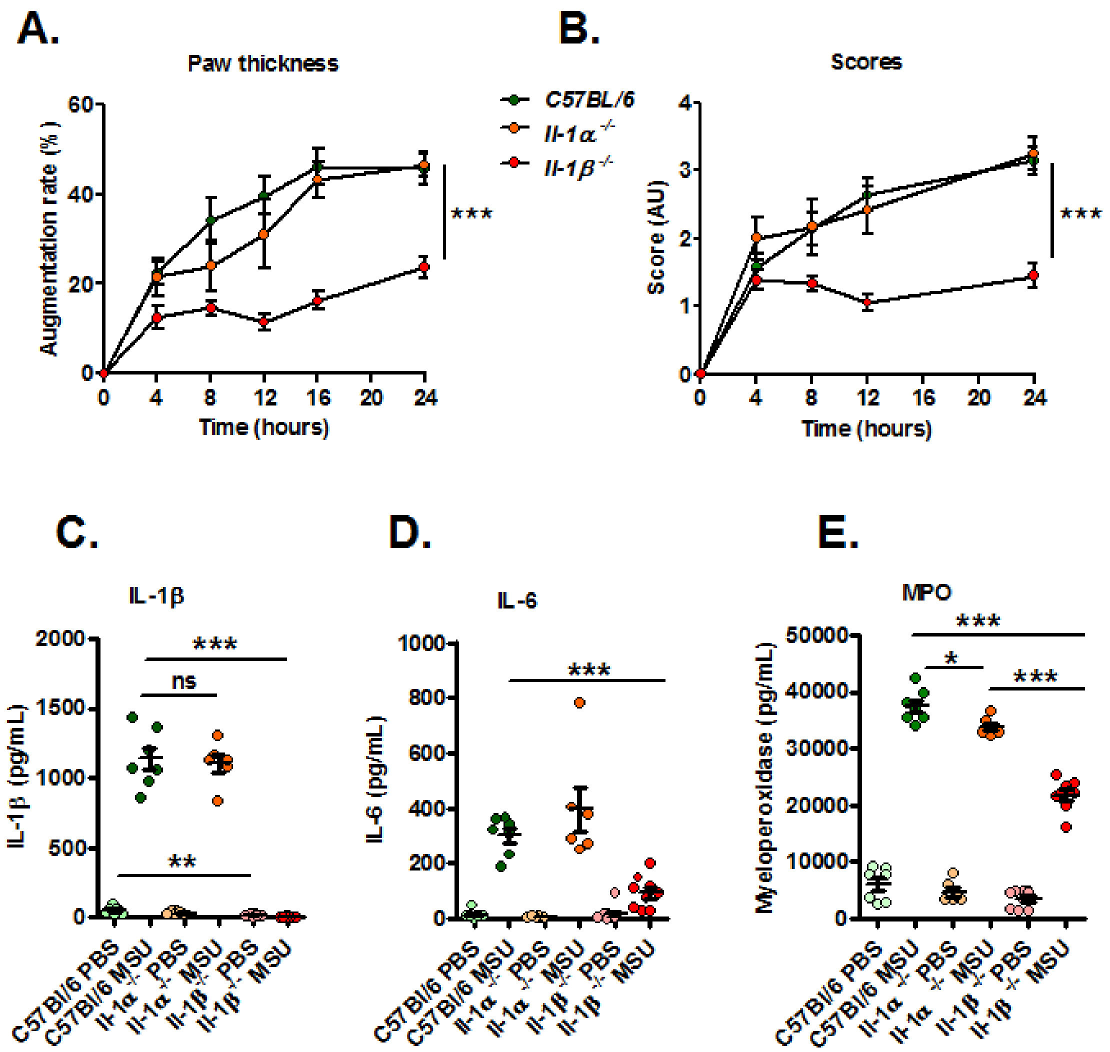

Supplm. Fig. S4

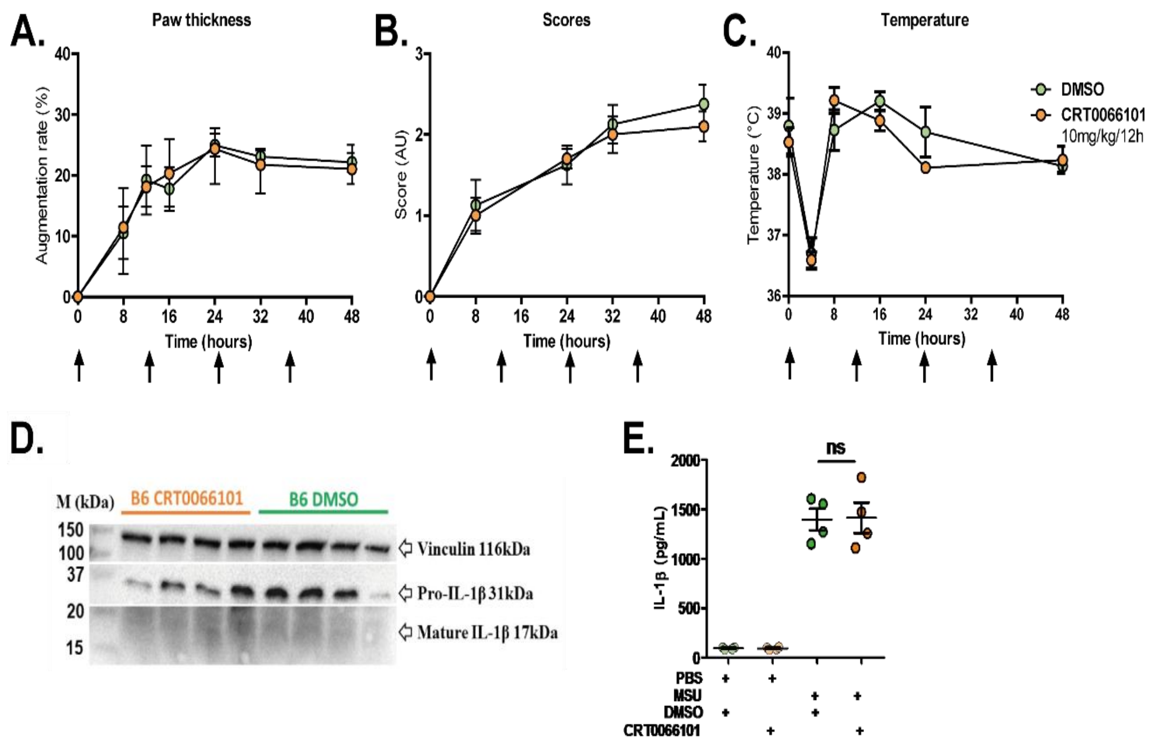

Supplm. Fig. S5

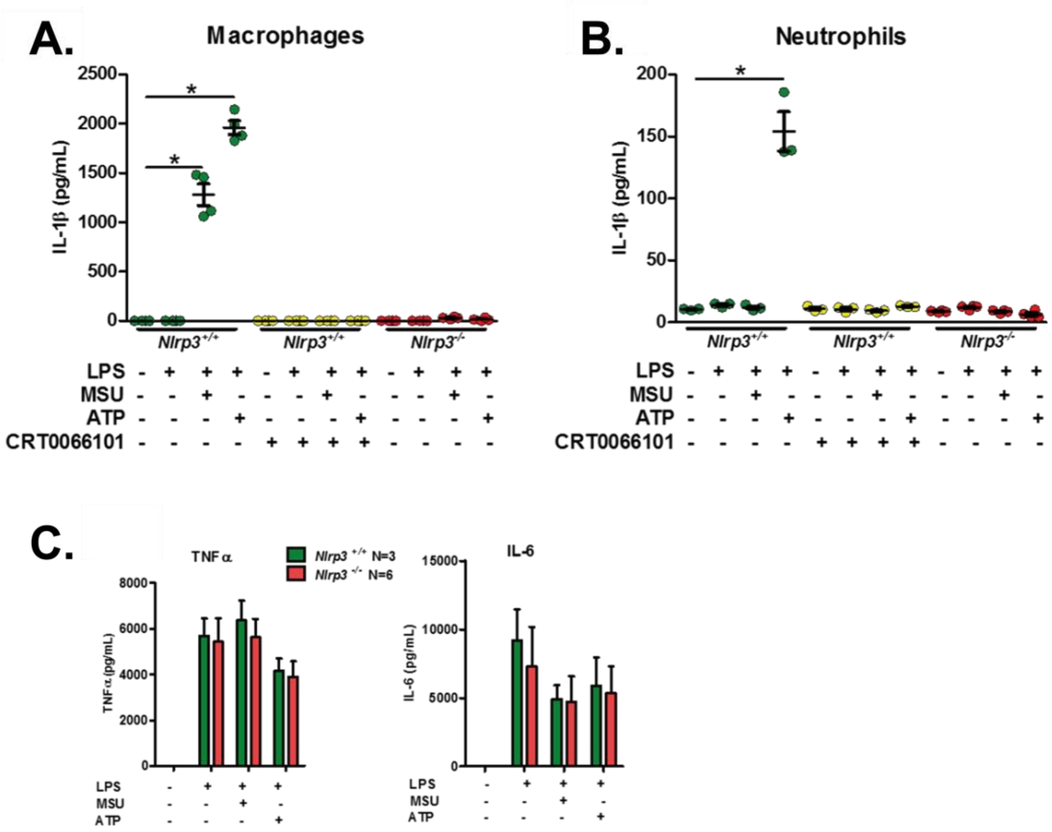

Supplm. Fig. S6

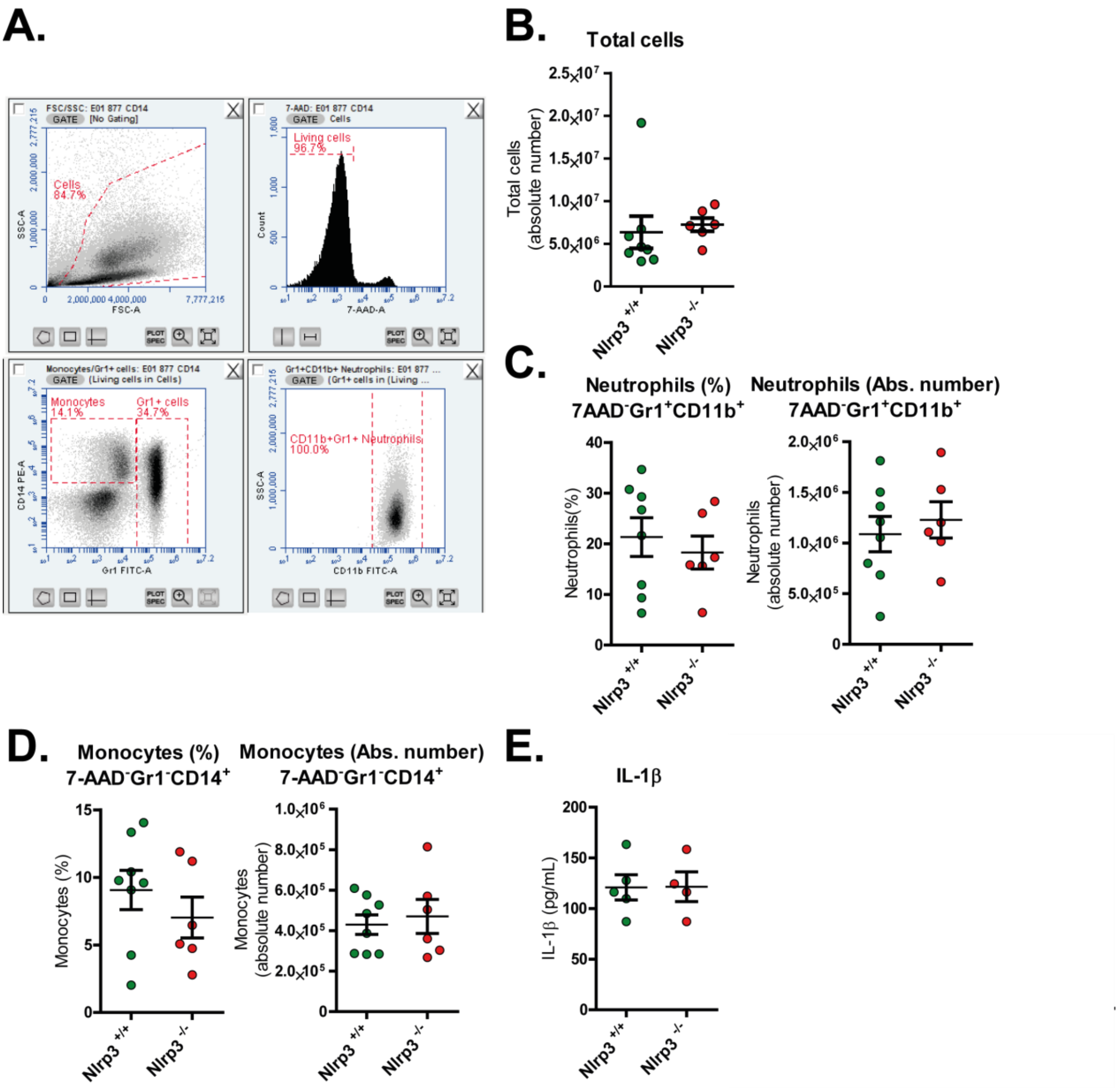

Supplm. Fig. S7

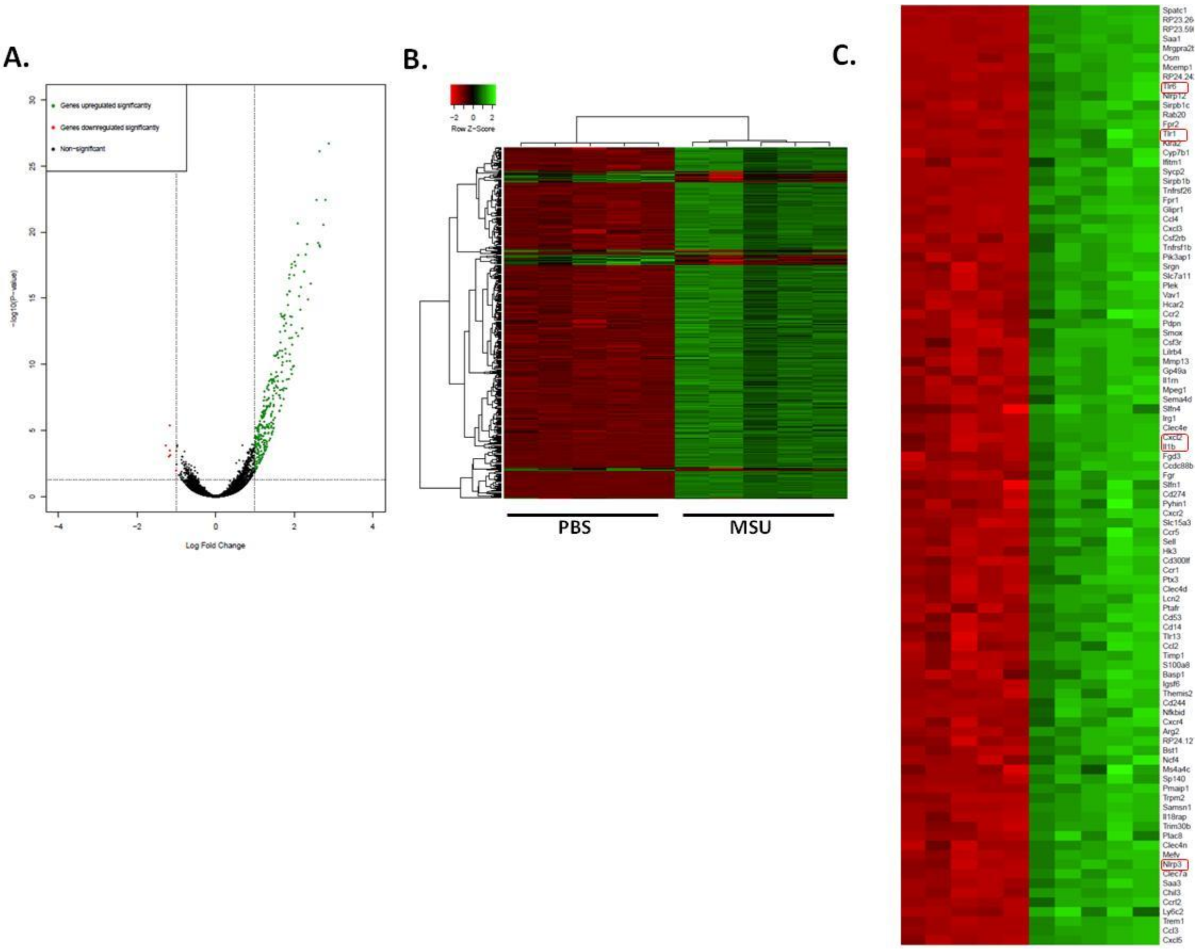

Supplm. Fig. S7 (continued)

**D.**

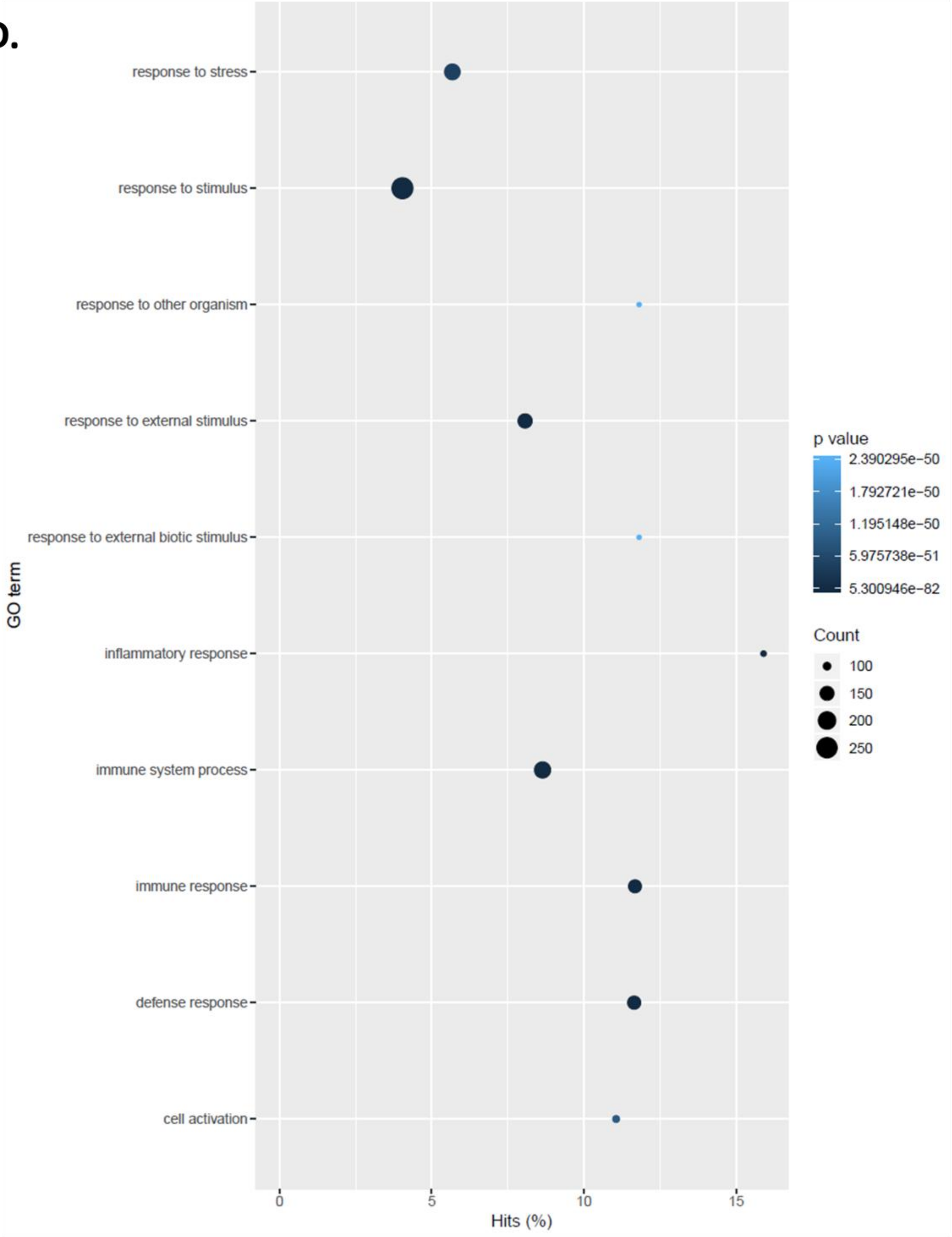

Supplm. Fig. S8

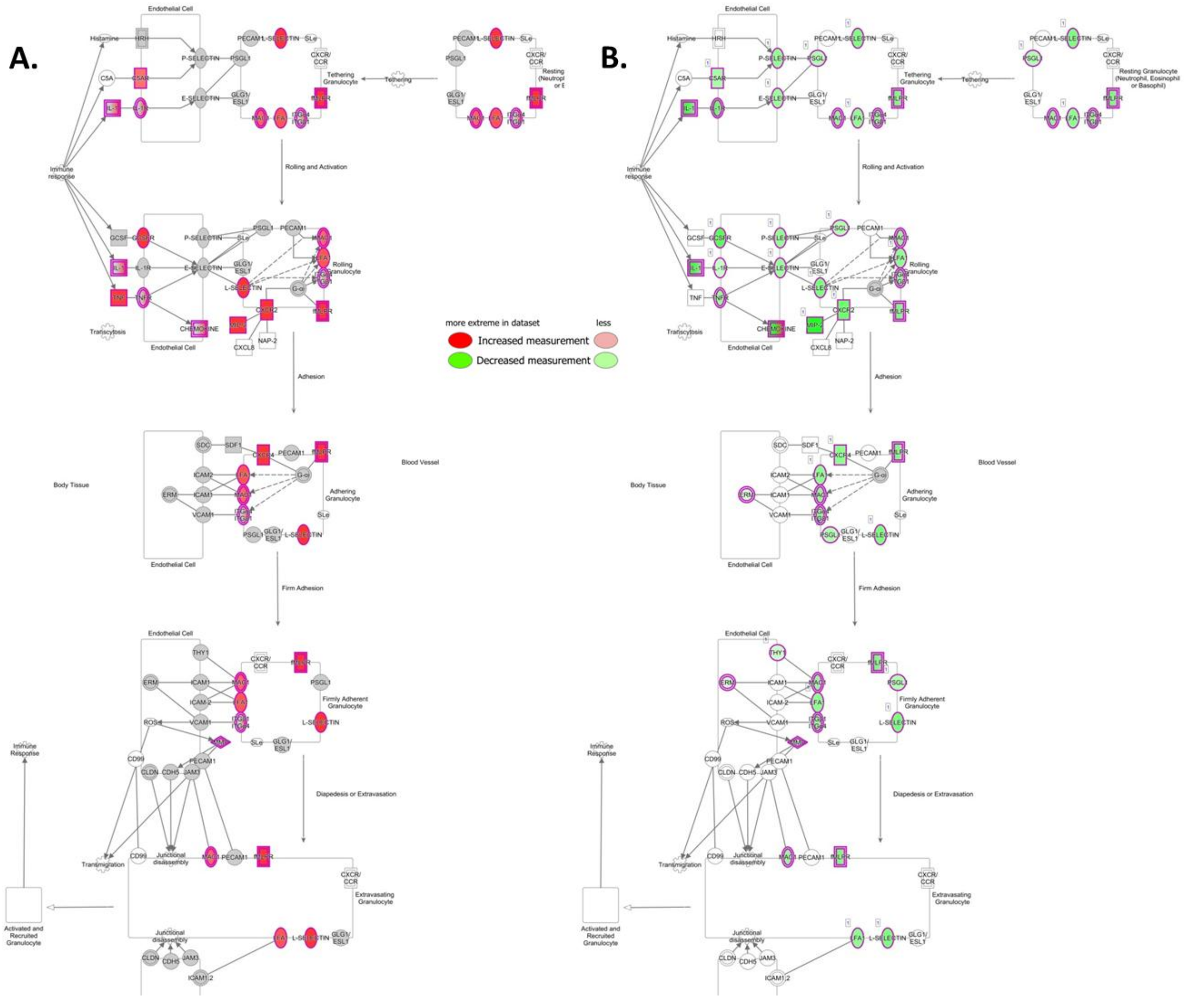

Supplm. Fig. S9

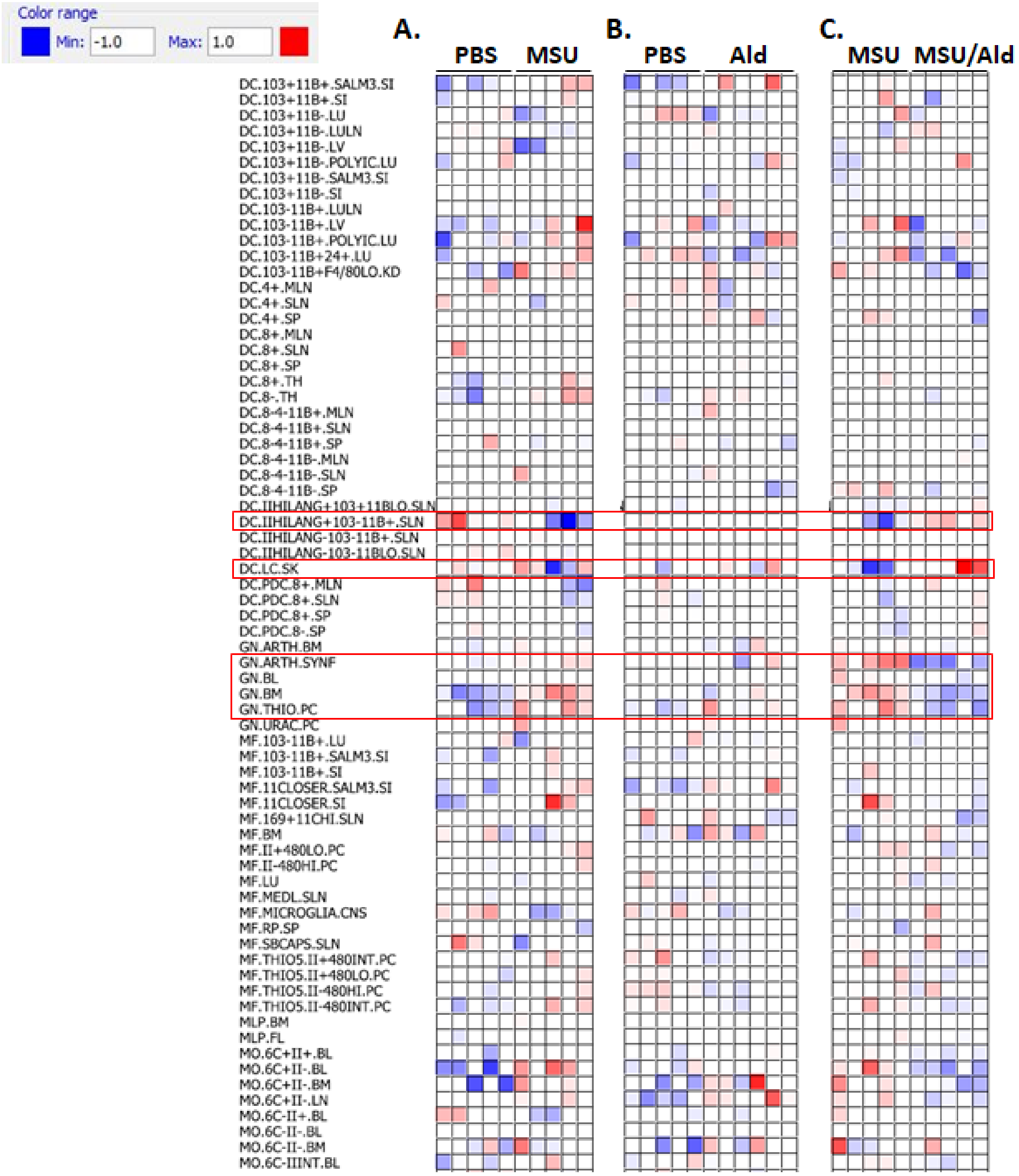

Supplm. Fig. S9 (continued)

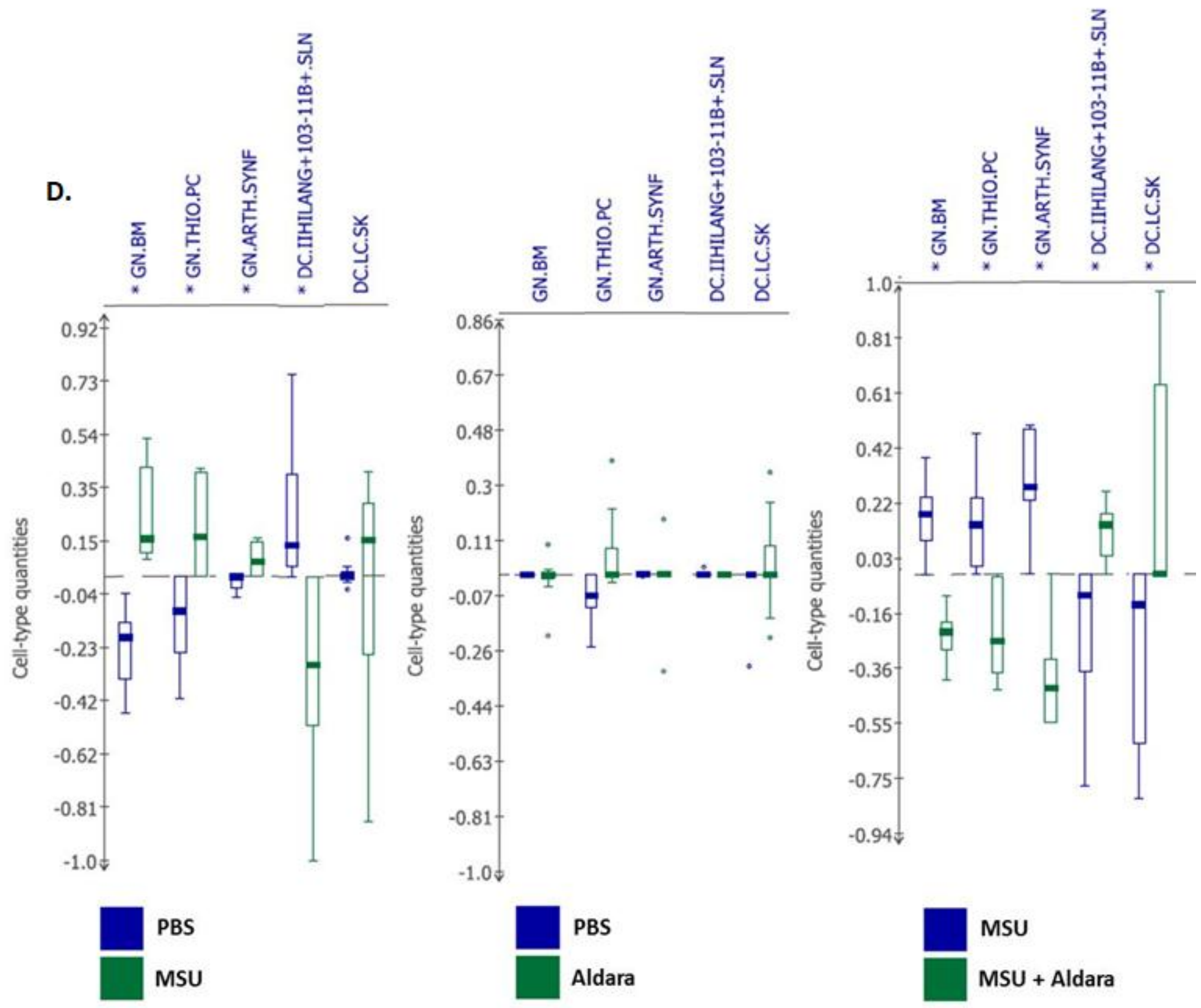

Supplm. Fig. S10

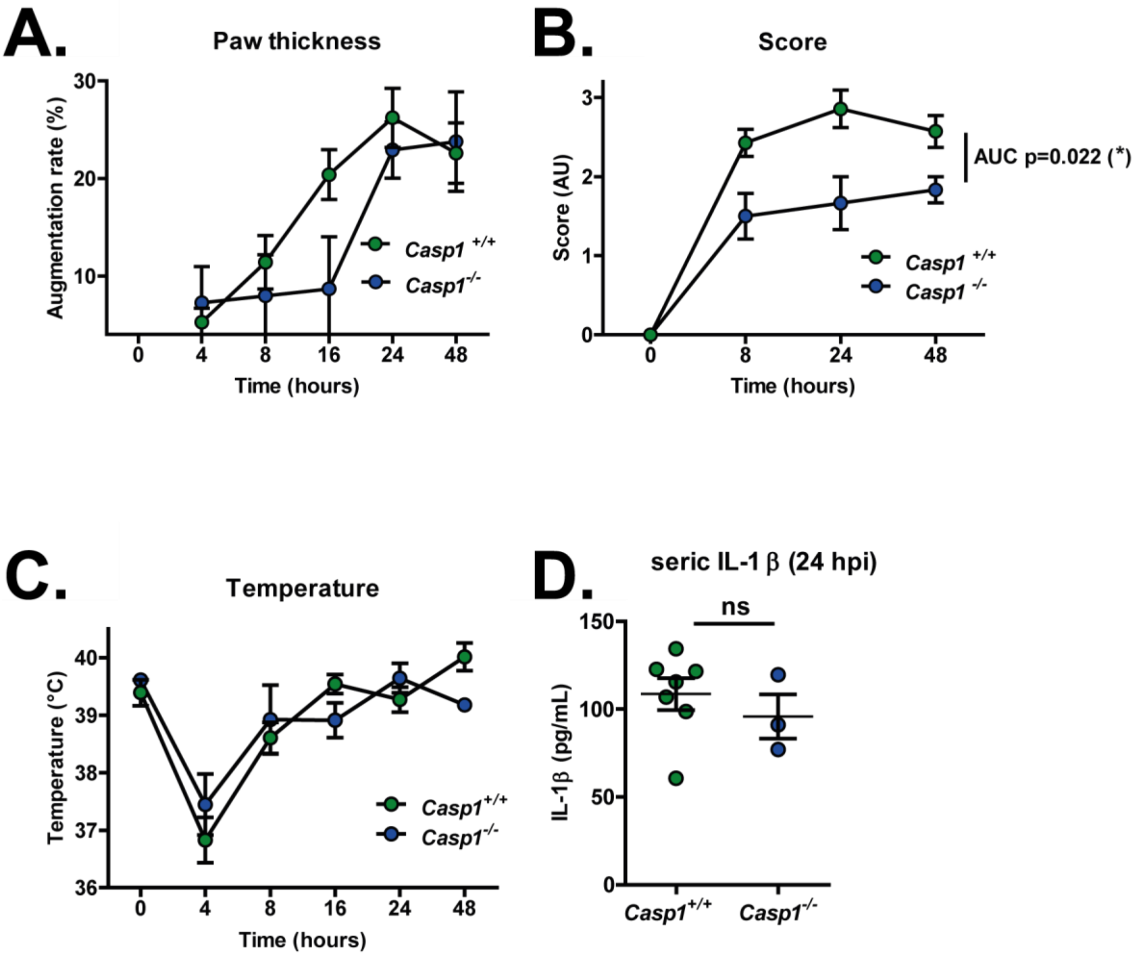

Supplm. Fig. S11

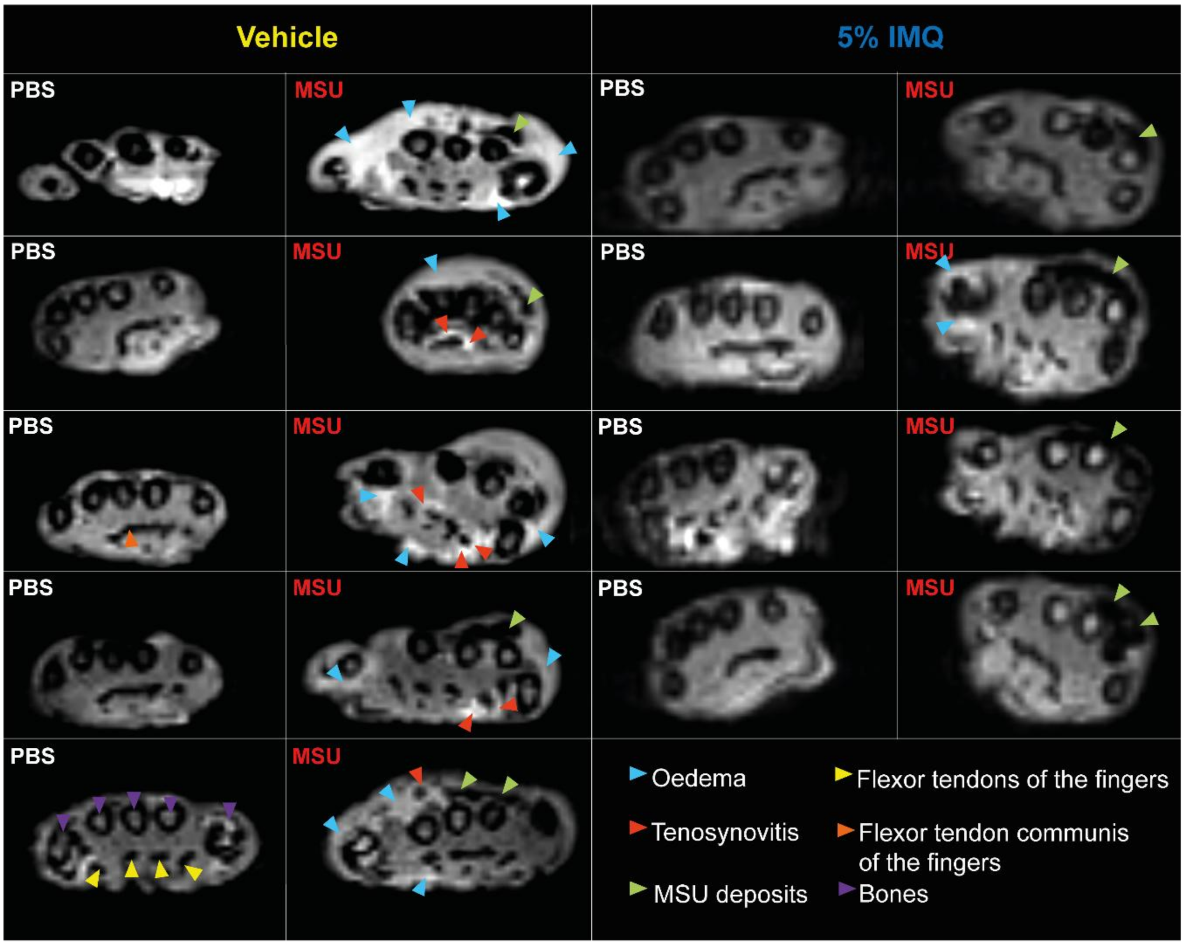

Supplm. Fig. S12

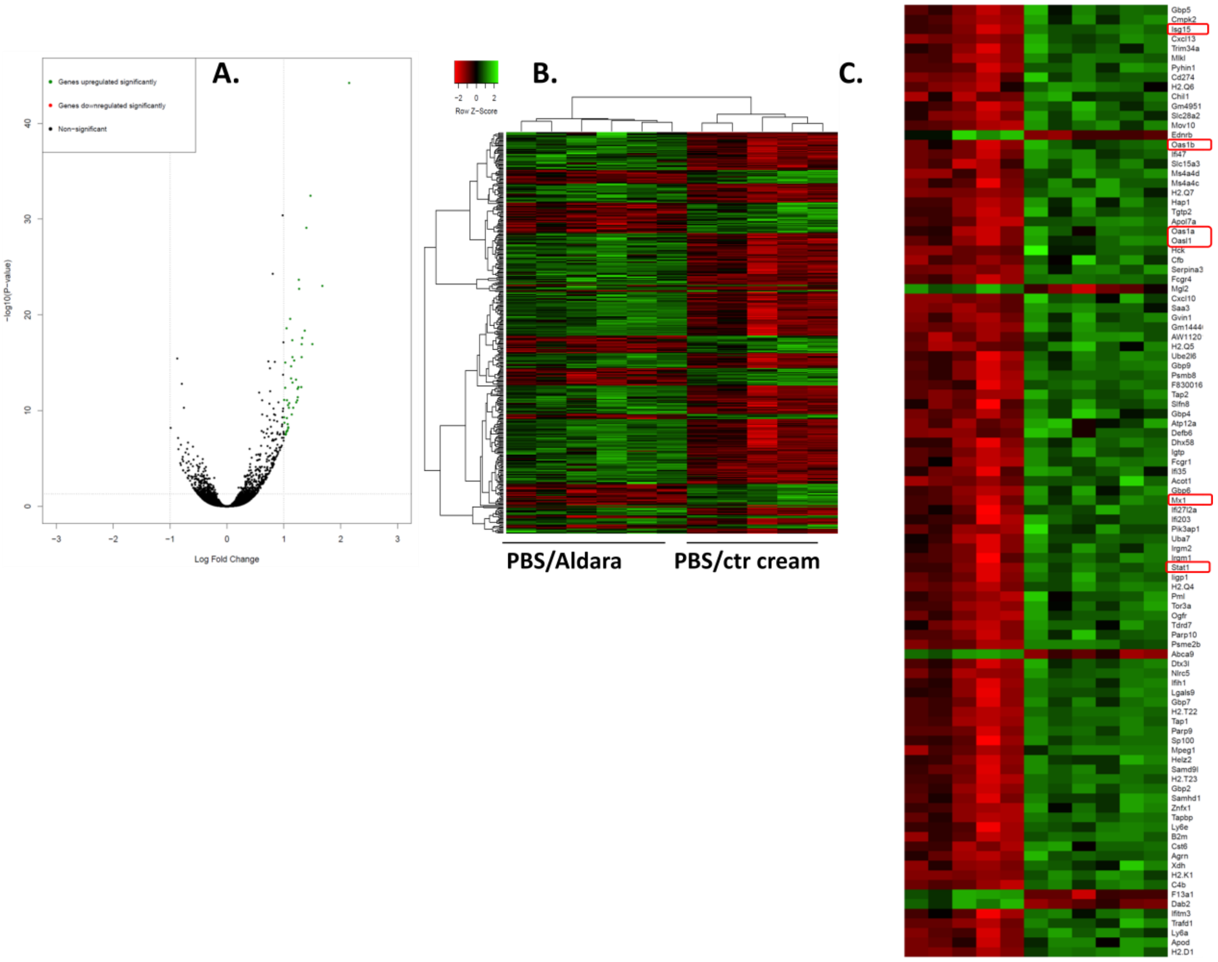

Supplm. Fig. S12 (continued)

D.

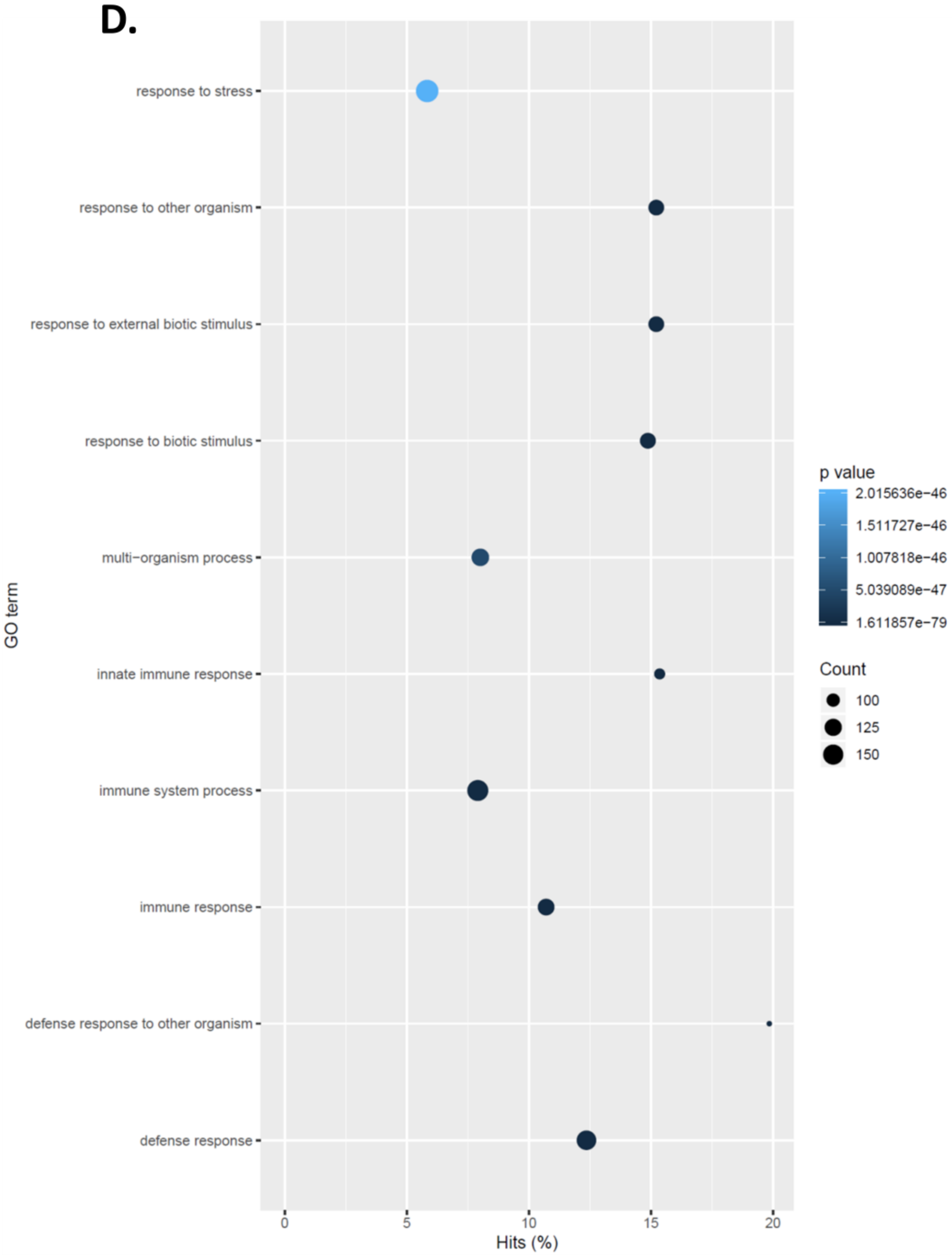

Supplm. Fig. S13

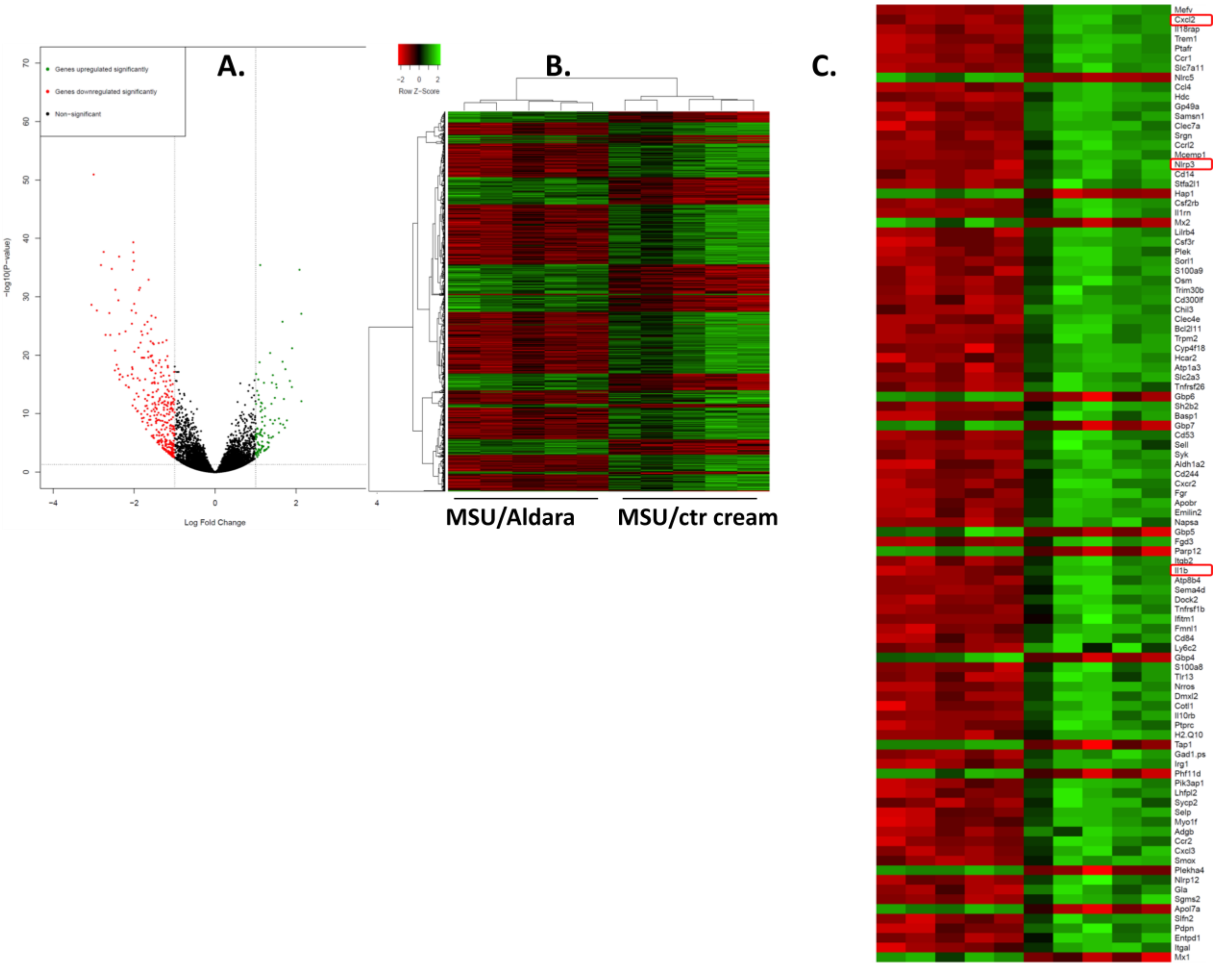

Supplm. Fig. S13 (continued)

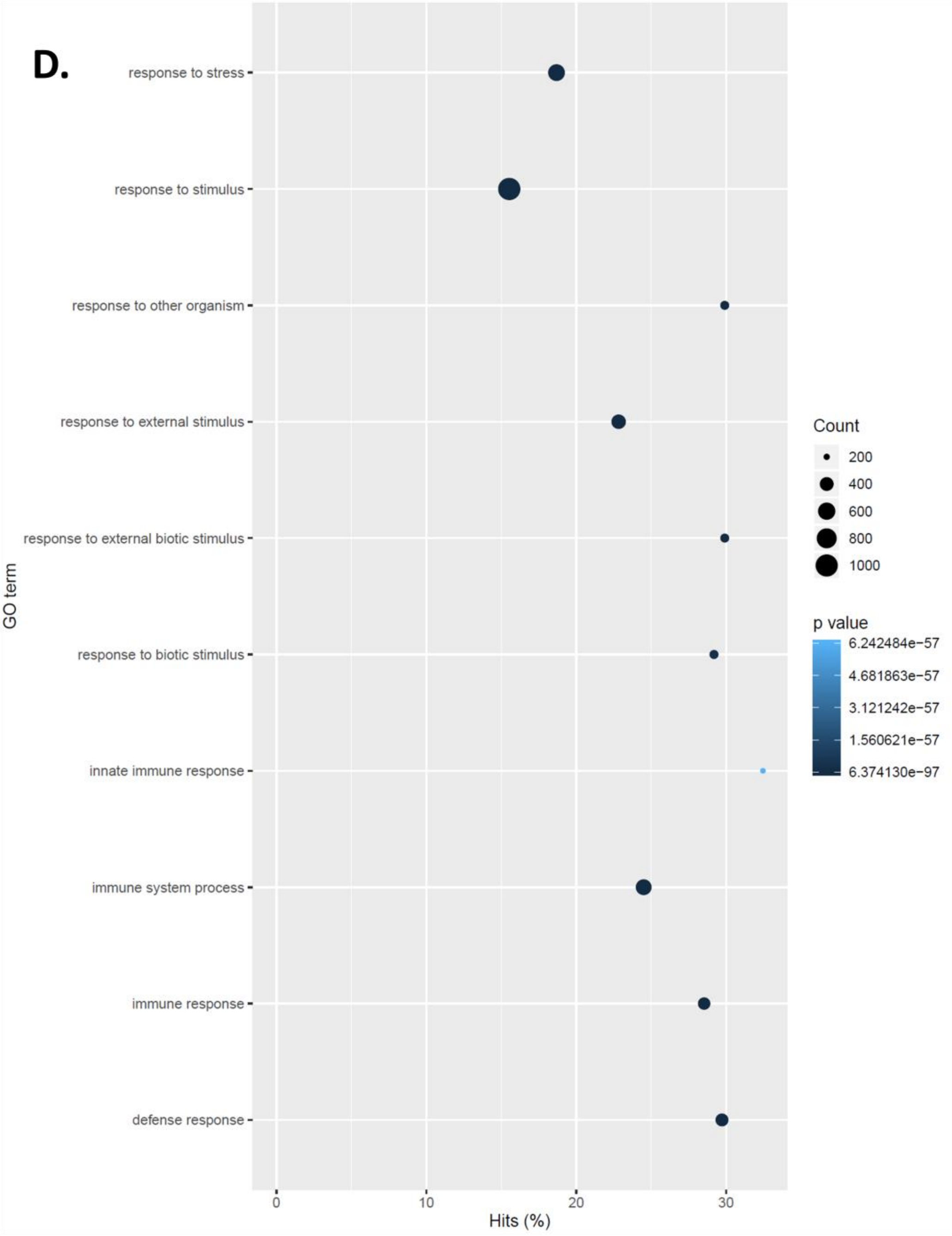

Supplm. Fig. S14

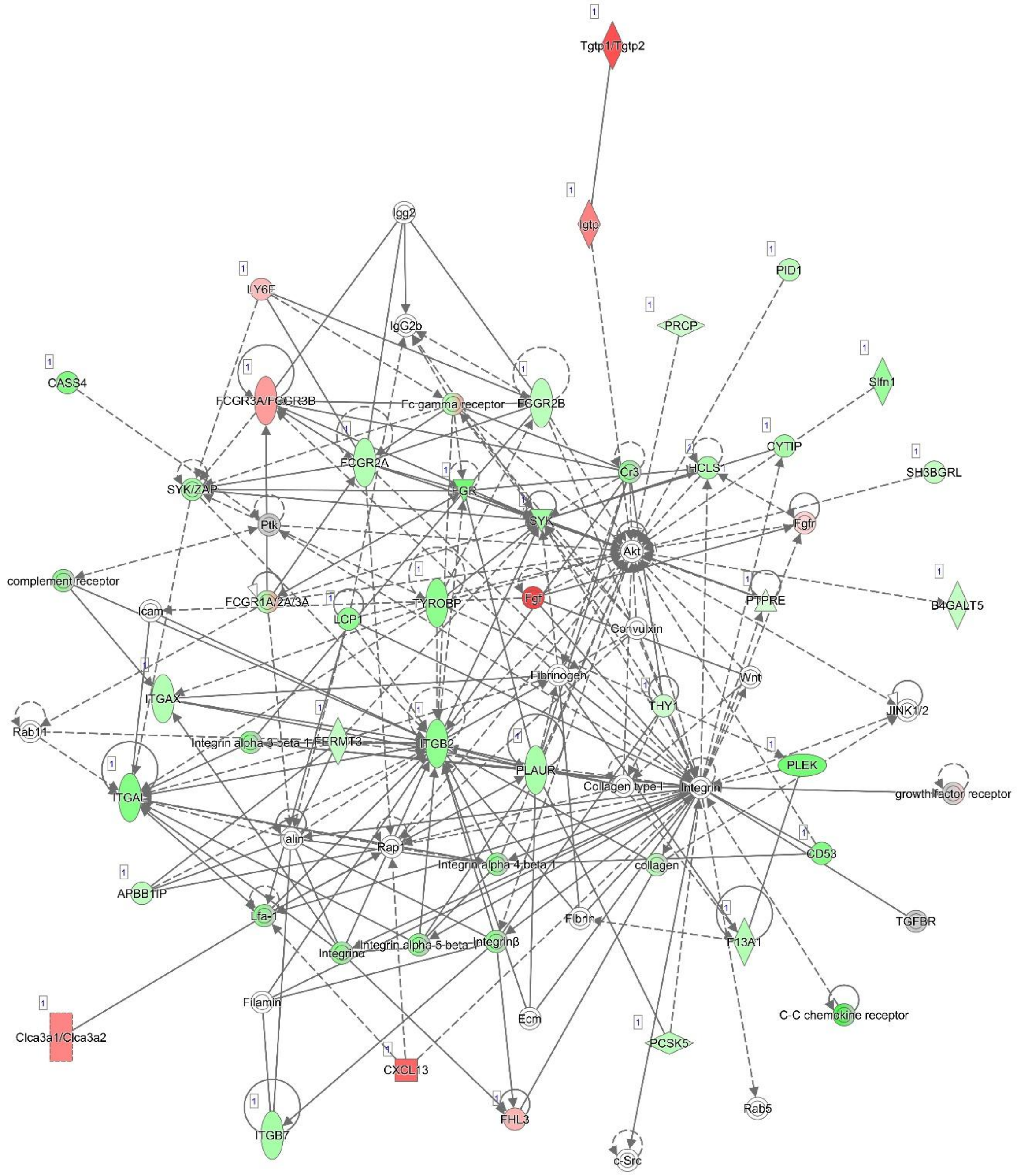


© 2000-2019 QIAGEN. All rights reserved.
